# Metavinculin mediates differentiation cues into myogenic gene expression

**DOI:** 10.1101/2024.02.21.578728

**Authors:** Abinayaselvi Murugan, Nandeesh Bevinahalli Nanjegowda, Mathivanan Jothi

**Affiliations:** Laboratory of Molecular Therapeutics, Department of Human Genetics, National Institute of Mental Health and Neurosciences (NIMHANS), Bengaluru, Karnataka, India; Department of Neuropathology, National Institute of Mental Health and Neurosciences (NIMHANS), Bengaluru, Karnataka, India

**Author notes:** Correspondence (M.J.).

**Keywords:** Metavinculin, Costameres, Alternative splicing, Rbfox proteins, Wnt signaling pathway, Wnt7b, Myogenesis, Skeletal muscle

## Abstract

Isoform switching in deciphering differentiation signals into myogenic transcriptional output remains unexplored. Here, we report a distinct function of muscle-specific metavinculin isoform in switching canonical to non-canonical Wnt-pathways to establish skeletal muscle differentiation. Metavinculin expression is specifically associated with muscle differentiation, regeneration and absent in proliferating myoblasts. During differentiation, metavinculin-specific exon retention is facilitated by the direct binding of Rbfox1 to its downstream intron. Depletion of metavinculin impairs skeletal muscle differentiation, while its ectopic expression induces genes associated with differentiation even in proliferating myoblasts. Subsequent results revealed that pharmacological activation of canonical Wnt impedes muscle differentiation, whereas its inhibition stimulates differentiation. Canonical Wnt inhibition or ectopic non-canonical Wnt7b expression restores muscle differentiation programs in metavinculin-depleted cells. These results indicates a dynamic interplay between Rbfox1-generated metavinculin and Wnt-signaling pathways. This advancement sheds light on the intricate molecular mechanisms underlying skeletal muscle differentiation and suggests potential therapeutic targets for muscle-related disorders.

## INTRODUCTION

Skeletal muscle differentiation is established by differentiation cues through a cascade of cellular events, which ultimately converge to alter the epigenome for differentiation-specific transcriptional output.^1,2,11,3–10^ It is tightly controlled by transcriptional factors like Myod1, Myf5, Myog, and Mrf4, collectively known as myocyte regulatory factors (MRFs) and myocyte enhancing factors (MEFs).^12,13^ Additionally, splicing factors are also involved in establishing myogenic gene expression as muscle is one of the human tissues with high differentially expressed alternative exons.^14–16^ Several proteins, switch ubiquitous to muscle-specific isoforms by distinct gene expression or alternative splicing that enable muscle cells adaptions in physiological conditions.^17–21^

Isoform switching is mainly observed in cytoskeletal proteins though it reported in other proteins.^18,19,22–25^ Cytoskeletal complex actively and constantly supports contracting muscle cells by modifying cellular response when they adhere to the extracellular matrix or contact with other cells.^26^ The cytoskeletal dynamics are influenced by various cell signaling pathways and vice versa during differentiation.^27–29^ Particularly, Serum responsive factor (SRF) function is regulated during muscle differentiation by cytoskeletal and sarcomeric dynamics through Myocardin related transcription factor (MRTF) and Striated muscle activator of Rho signaling.^30–32^ Further, altered cytoskeleton activates TGF-β signaling to enhance myofibroblasts conversion in β-actin knockout mouse embryonic fibroblasts.^33^ However, how isoform switching in cytoskeletal proteins contribute myogenic signaling during differentiation remains unexplored. Vinculin is a ubiquitous cytoskeletal scaffold protein localized to focal adhesions and adherent junctions.^34^ It involved in cell morphology control, integrin clustering, motility, force transduction and adhesion strength.^34^ In addition, it may also have regulatory or signaling roles which largely unexplored.^35^ Vinculin knockout embryos are not viable and showed retarded somites and limbs in addition to severe cardiac and neuronal defects.^36^ Metavinculin, a longer muscle-specific splice variant of vinculin generated by alternative splicing of exon 19 that encodes 68-amino acid residues, is co-localized with vinculin at the costameres, sarcolemma, I band in the sarcomere and intercalated discs.^37–42^ Its expression was reported in smooth, cardiac, skeletal muscles and platelets.^43–47^ Metavinculin loss or point mutations at its unique exon is linked with dilated and hypertrophic cardiomyopathies.^44,48–53^ Cholinergic receptor stimulation recruits metavinculin into the integrin binding complex in airway smooth muscle.^54^ Several proteins activate vinculin by disrupting its auto-inhibitory head and tail interaction and metavinculin forms heterodimer with activated vinculin.^44,54–57^ Upon activation vinculin binds with actin filaments (F-actin), facilitating vinculin dimerization for its bundling. Conversely, metavinculin binding to F-actin inhibits vinculin dimerization, thereby preventing actin bundling.^58–62^ Further, metavinculin-specific insert sequestered as a globular subdomain protruded from the F-actin surface also prevents its assembly.^59^ Contrastingly, metavinculin reduces F-actin bending persistence, increasing susceptibility to breakage.^63^ In addition, it also increases cell adhesion complex stability by altered force propagation with enhanced shear force resistance.^45^ These studies linked metavinculin with actin organization and force transduction and the significance of its unique expression in muscle cells left unexplored from its discovery.^40,58–63^

Here, we have observed differentiation-specific metavinculin isoform expression in skeletal muscle cells. Several splicing factors regulate alternative splicing in muscle cells, however, the factor/s responsible for metavinculin generation has remained unknown.^40,41,64–67^ Here, we report Rbfox1 regulates metavinculin splicing through direct binding at its downstream intronic region during muscle differentiation. Further, depletion, ectopic expression and transcriptomic analysis showed its importance of metavinculin in executing skeletal muscle differentiation. Subsequently, muscle differentiation defects observed in metavinculin-depleted cells is restored by either canonical Wnt-inhibition or non-canonical Wnt7b ectopic expression. Together, all our evidences suggest that metavinculin act as a critical mediator in switching canonical to non-canonical Wnt-signaling to establish skeletal muscle differentiation.

## RESULTS

### Metavinculin is induced during skeletal muscle differentiation and regeneration

Metavinculin expression was found in smooth and cardiac and skeletal muscles.^37–39,47,68^ Here, we verified the sub-stoichiometric expression of the 124 kDa metavinculin isoform in heart, intestine, and tibialis anterior (TA) muscle tissues from C57BL/6J mice (Figure 1A). We have collected 50% confluent C2C12 (Mouse), L6 (Rat), and HSMM (Human) cells grown in high serum containing media as GM and after reaching 80% confluency they are switched to low serum (DM) media to induce differentiation. We confirmed the cell cycle inhibitor Cdkn1a induction in all muscle cells cultured in differentiation media as growth arrest is a prerequisite for differentiation (Figure S1A).^12,69^ In vitro skeletal muscle differentiation was confirmed by the expression of early myogenic markers Myod1, Myog, and late marker Myh (Myosin heavy chain) at RNA (Figure S1A) and protein levels (Figure 1B). Further, we have also estimated the fusion index in these cells using Myh immunofluorescence images (Figure S1B). We have found metavinculin protein expression in C2C12, L6, and HSMM cells cultured under differentiation-promoting condition not in the growth-promoting conditions whereas vinculin expression remains unaltered (Figure 1B). Similar results are observed at transcript levels (Figure 1C). Further, we could detect metavinculin protein expression as early as twelve hours of differentiation induction and it is increased with incubation time (Figure 1D). Thus, these results showed metavinculin isoform is specifically expressed during skeletal muscle differentiation (Figure 1B-D).

**Figure 1:**
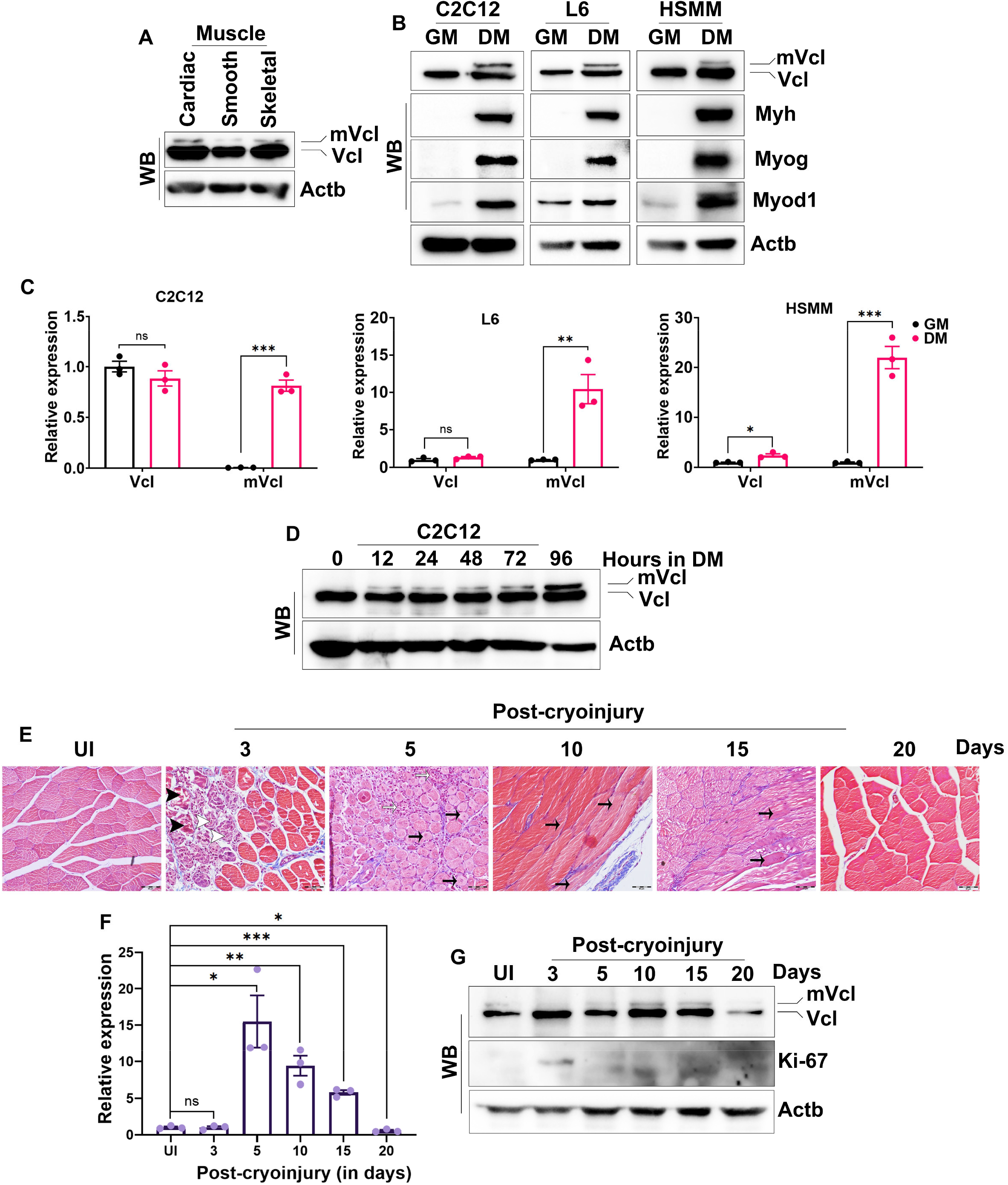
Metavinculin isoform expression during skeletal muscle differentiation and regeneration, See also in Figure S1, Table S1. (A) Western blot showing metavinculin and vinculin expression in cardiac, smooth and skeletal muscle tissues. (B) Expression of vinculin isoforms and myogenic genes Myod1, Myog and Myh expression in C2C12, L6 and HSMM myoblasts grown under growth (GM) and differentiation (DM) media. (C) RT-qPCR showing the relative expression of vinculin and metavinculin transcripts in various myoblasts cultured under growth and differentiation conditions. (D) Western blot showing the vinculin isoform expression at indicated time points during differentiation. (E) Regenerating muscle tissue sections showing myonecrosis (Black arrow head), myophagocytosis (White arrow head), inflammation (White arrow) and regenerating fibers (Black arrow) in different days of post cryoinjury. Scale bar 50 μM. (F) Metavinculin mRNA expression during in vivo muscle regeneration is shown in a bar graph. (G) Western blot showing vinculin isoforms and marker of cell proliferation Ki-67 expression in regenerating muscle tissues. Actb acted as loading control in western blot images. Data are shown as mean value ±SEM, n=3. p value * ≤0.05; **≤0.01; ***≤0.001; ns-not significant.

Mouse muscle injury models are well-established and extensively used systems to study muscle regeneration. Here, we have utilized in vivo mouse cryoinjury model to analyze the metavinculin expression during skeletal muscle regeneration.^70^ Specifically, we have created cryoinjury in the hind limb TA muscle of three-month-old male C57BL/6 mice using a metal rod precooled in dry ice. We have obtained the regenerating tissues and uninjured contralateral TA muscle as control from the same animal at different post-injury time points. First, muscle regeneration process are evaluated by histopathological examination using Hematoxylin and eosin (H&E) (Data not shown) and Masson’s trichrome staining (MTS). We have observed typical histological features of muscle regeneration after muscle injury (Figure 1E). Specifically, the three days post-injured muscle showed active myonecrosis and myophagocytosis in the injured area. The muscle regeneration appeared from the viable zone on day five post-injured muscle that showed resolving myonecrosis and inflammation. We have noticed more regenerating fibers infiltrate the injured area with focal inflammation on day ten and these fibers completely occupied the injured area on day fifteen. At twenty days post-injury the damaged muscle completely regenerated (Figure 1E). RT-qPCR showed induction of metavinculin transcript at day five and decreased gradually until completion of regeneration (Figure 1F). Similarly, metavinculin protein expression is increased in ten days post-injured muscle and decreased in later time points (Figure 1G). Strikingly, we have noticed a transient metavinculin protein reduction in three days post-injured muscle compared to uninjured and marker of proliferation Ki-67 also appeared at this time point (Figure 1G). This further supported the negative correlation of metavinculin expression with proliferation. Hence, our in vitro and in vivo experiments revealed metavinculin expresses in differentiating not in proliferating cells (Figure 1B-1G).

### Splicing regulator Rbfox1 retains metavinculin-specific exon during skeletal muscle differentiation

Muscle-specific metavinculin expression was identified more than thirty years ago however the splicing factor regulates vinculin alternative splicing is not identified yet.^40,41^ Although several splicing factors include CELF, MBNL, Fox, and PTB families regulate alternative splicing in the muscle cells,^64–67^ they have very low tissue specificity score except Rbfox family (Table S2). Notably, skeletal muscle-specific enhanced expression was observed only for Rbfox1 with the tissue specificity Tau index 0.82 (Table S2). Here, we have found differentiation-associated induction of Rbfox1, no change in Rbfox2, and variable expression of Rbfox3 mRNAs in C2C12 and L6 cells (Figure 2A and 2B). Rbfox1 and Rbfox2 are associated with skeletal muscle and Rbfox3 with neuronal development.^18,19,66,71^ Therefore, we have focused on Rbfox1 and Rbfox2 in metavinculin generation and excluded Rbfox3.

**Figure 2:**
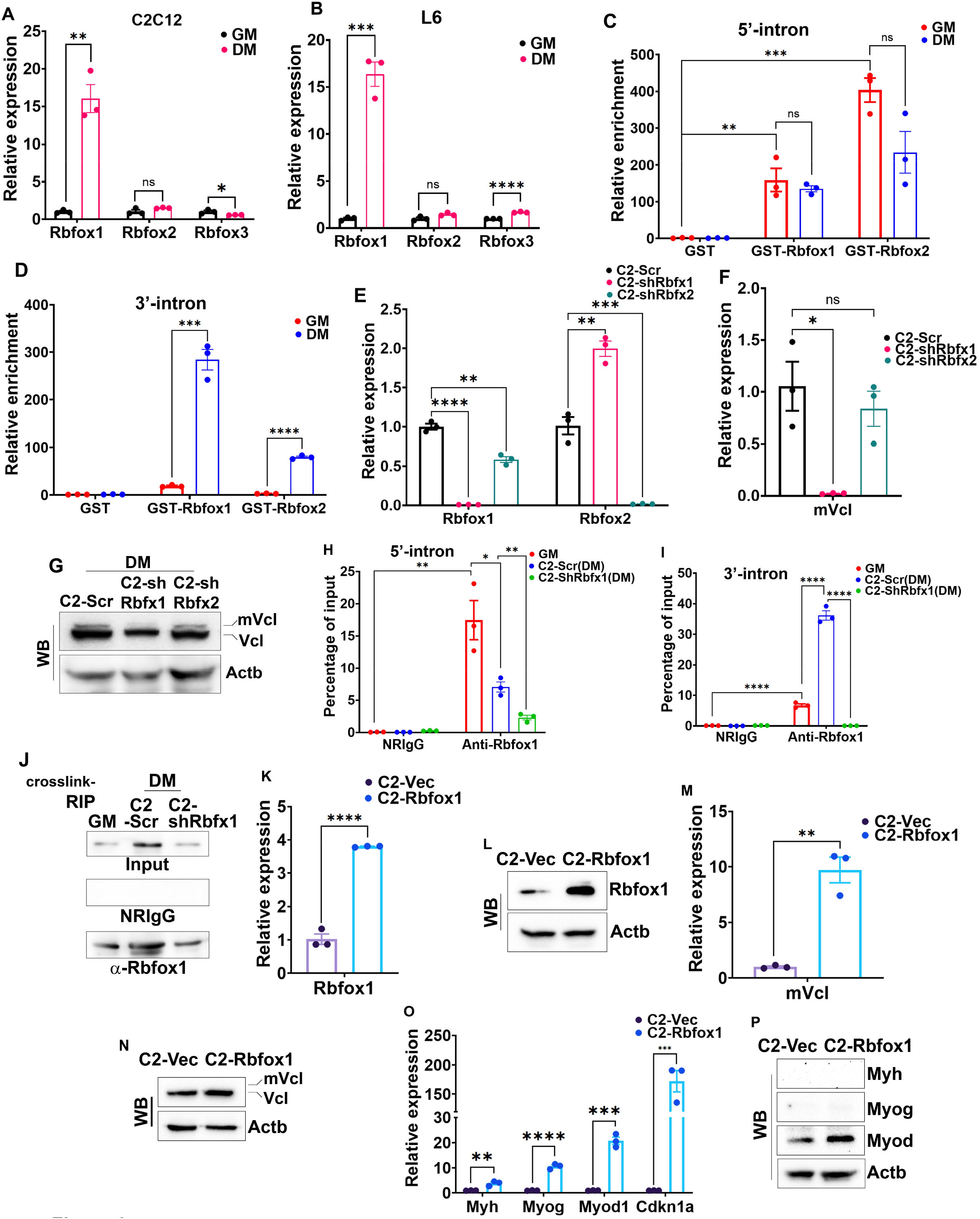
Rbfox1 expression regulates metavinculin splicing during skeletal muscle differentiation, See also in Figure S2. (A and B) Relative expression of Rbfox family members during in vitro differentiation of C2C12 (A) and L6 (B) myoblasts are shown in RT-qPCR. (C and D) In vitro GST-pulldown assay using total RNAs isolated from C2C12 cells grown in GM and DM media showing relative enrichment of Rbfox1 and Rbfox2 with 5’ (C) and 3’ (D) intron flanking metavinculin-specific exon. (E) Rbfox1 and Rbfox2 mRNA expression in Rbfox depleted C2-shrbfx1, C2-shRbfx2 and control C2-Scr cells shown in a bar graph. (F and G) Metavinculin expression in Rbfox depleted cells shown by RT-qPCR (F) and western blot analysis (G). (H and I) Crosslink-RIP showing the enrichment of Rbfox1 at upstream (H) and downstream (I) metavinculin flanking intron in C2-Scr and C2-shRbfx1 cells grown under mentioned culturing conditions. (J) Western blot image showing Rbfox1 level in Anti-Rbfox1 or NRIgG control immunoprecipitate along with input samples. (K-N) Rbfox1 and metavinculin expressions are shown in C2-Vec and C2-Rbfox1 cells having exogenous vector or Rbfox1 cDNA at mRNA (K and M) and protein levels (L and N). (O) Induction of muscle differentiation-associated genes in growth conditions by ectopic Rbfox1 is shown in a bar graph. (P) Western blot image of myogenic proteins in C2-Vec and C2-Rbfox1 cells. Actb served as loading control in western blot images. Data are shown as mean value ±SEM, n=3. p value * ≤0.05; **≤0.01; ***≤0.001; ****≤0.0001; ns-not significant.

Rbfox family recognizes auxiliary cis element 5’-GCAUG on the target RNA and regulates exon inclusion or exclusion depending on its location.^66^ We have analyzed the putative Rbfox binding element within the 1kb intronic region that flanks metavinculin-specific exon using RBPmap Ver. 1.2 and found two potential Rbfox binding sites in the upstream and eight in the downstream intron region (Figure S2A). To validate Rbfox1/2 binding in metavinculin-flanking introns we have utilized GST-pulldown assay using recombinant GST-Rbfox1, GST-Rbfox2, and control GST (Figure S2B). Here, GST-tagged proteins are incubated with the total RNAs isolated from C2C12 cells grown in GM and DM media. The bound RNAs are analyzed by RT-qPCR using specific primers directs putative Rbfox binding sites (Table S1). We have noted GST-Rbfox1 and GST-Rbfox2 binding to the 5’-intronic flank in both GM and DM conditions (Figure 2C). Further, we have observed the differentiation-specific enrichment at the 3’-intronic flank by both the proteins, however, GST-Rbfox1 enriched more than GST-Rbfox2 (Figure 2D). To identify the Rbfox protein that is responsible for metavinculin generation we have created stable C2-shRbfx1, C2-shRbfx2, and control C2-Scr cells using lentivirus-mediated transduction shRNAs targeting Rbfox1, Rbfox2 and scramble shRNA (Key resources table). Rbfoxs depletion in these cells are confirmed by RT-qPCR analysis (Figure 2E). Surprisingly, we have observed that Rbfox1-depletion significantly increased the Rbfox2 mRNA expression whereas Rbfox2-depletion reduced Rbfox1 mRNA expression (Figure 2E). Metavinculin mRNA and protein levels are reduced in C2-shRbfx1 cells compared to C2-shRbfx2 and scramble C2-Scr (Figure 2F and 2G). Further, we have found that Rbfox1-depletion completely inhibits the expression of early and late myogenic genes at RNA and protein levels (Figure S2C and S2D). The differentiation defect in these cells is also noted by reduced myotube formation and fusion index (Figure S2E and S2F). Although Rbfox2-depletion inhibited skeletal muscle differentiation it is not as effective as Rbfox1-depletion (Figure S2C-S2F). However, Rbfox2-depletion affects skeletal muscle differentiation while retaining metavinculin expression (Figure 2F and 2G, S2C-S2F). These results suggest that Rbfox1 is needed for metavinculin expression during muscle differentiation (Figure 2E-2G).

To further confirm Rbfox1 in metavinculin generation, we have performed crosslink-RNA immunoprecipitation (crosslink-RIP). Here, we have used the C2-Scr and C2-shRbfx1 cells cultured in DM conditions and C2-Scr in GM as an additional control to analyze Rbfox1 bound RNAs using anti-Rbfox1 and isotype control antibodies. RT-qPCR analysis of crosslink-RIP RNAs showed metavinculin flanking intron regions were enriched only with anti-Rbfox1 not in the NRIgG (Figure 2H and 2I). Specifically, metavinculin 5’-intron is enriched in anti-Rbfox1 crosslink-RIP under growth conditions and it is decreased during differentiation conditions (Figure 2H). In contrast, 3’-intron is enriched in RIP under differentiation conditions compared to the growth (Figure 2I). The presence of Rbfox1 is confirmed in input and immunoprecipitate by western blot analysis (Figure 2J). As expected, Rbfox1-depletion abrogates these enrichments (Figure 2H and 2I). These results further confirmed that Rbfox1 specifically switches its binding from the 5’-metavinculin intronic region to the 3’-intronic region during skeletal muscle differentiation thereby retains metavinculin-specific exon (Figure 2H and 2I).

We have generated stable C2-Rbfox1 and C2-Vec cells that ectopically express Rbfox1 cDNA or vector in C2C12 cells by retroviral transduction to further validate Rbfox1 in metavinculin generation. Ectopic expression of Rbfox1 is confirmed in RT-qPCR (Figure 2K) and western blot analysis (Figure 2L). Rbfox1-mediated Mef2d splicing pattern was also observed in C2-Rbfox1 cells as reported earlier (Figure S2G).^19^ Ectopic Rbfox1 induced metavinculin mRNA compared to vector control, however, it is not reflected in the protein level (Figure 2M and 2N). The myogenic differentiation genes are also induced only at RNA levels in C2-Rbfox1 cells compared to C2-Vec (Figure 2O and 2P). Nevertheless, these results also support the involvement of Rbfox1 in metavinculin exon inclusion in growth-promoting conditions. Collectively, these evidences strongly suggest that Rbfox1 occupy downstream flanking intronic region to include metavinculin-specific exon during skeletal muscle differentiation (Figure 2).

### Metavinculin expression is required for skeletal muscle cell differentiation

Metavinculin expression found only in differentiating myoblasts (Figure 1).^37,43,45^ We have designed and cloned two shRNAs (shmVcl1 and shmVcl2) targeting metavinculin-specific exon region and generated cell lines C2-shmVcl1 and C2-shmVcl2 that stably express these shRNAs and scramble control C2-Scr by lentiviral mediated transduction to identify differentiation-specific role of metavinculin. The effective metavinculin depletion is found by shmVcl2 compared to shmVcl1 shRNA at RNA and protein levels and vinculin levels are not altered by these shRNAs (Figure 3A and 3B). Therefore, we have used shmVcl2 shRNA to generate L6-shmVcl2 from L6 rat myoblast and scramble control L6-Scr cells as mentioned above. RT-qPCR results showed the inhibition of muscle-specific genes Myod1, Myog, Myh, and growth inhibitor Cdkn1a in metavinculin-depleted C2C12 and L6 cells cultured under DM media (Figure 3C and 3D). Similarly, differentiation-associated myogenic proteins are also repressed by metavinculin depletion in these cells (Figure 3E). The inhibition of myogenic gene activation by metavinculin depletion severely affected myotube formation as shown by Myh immunofluorescence and fusion index (Figure 3F). Myod1 occupancy to its target gene Myog and Myh promoters is also inhibited by metavinculin depletion in C2C12 cells (Figure S3A and S3B). These experiments revealed that metavinculin expression is necessary to induce myogenic gene expression during skeletal muscle differentiation (Figure 3C-3F). Therefore, we have verified its ability to induce these genes at growth-promoting conditions. For this we have transiently expressed GFP-tagged metavinculin in C2C12 and L6 myoblasts and analyzed for myogenic gene expression after 48 hours of post-transfection. The ectopic expression of metavinculin is confirmed using western blot analysis (Figure 3G). Metavinculin ectopic expression is sufficient to induce myogenic marker genes Myod, Myog, and Myh even at growth-promoting conditions in C2C12 and L6 cells (Figure 3G). These results revealed the importance of metavincluin in inducing myogenic gene expression.

**Figure 3:**
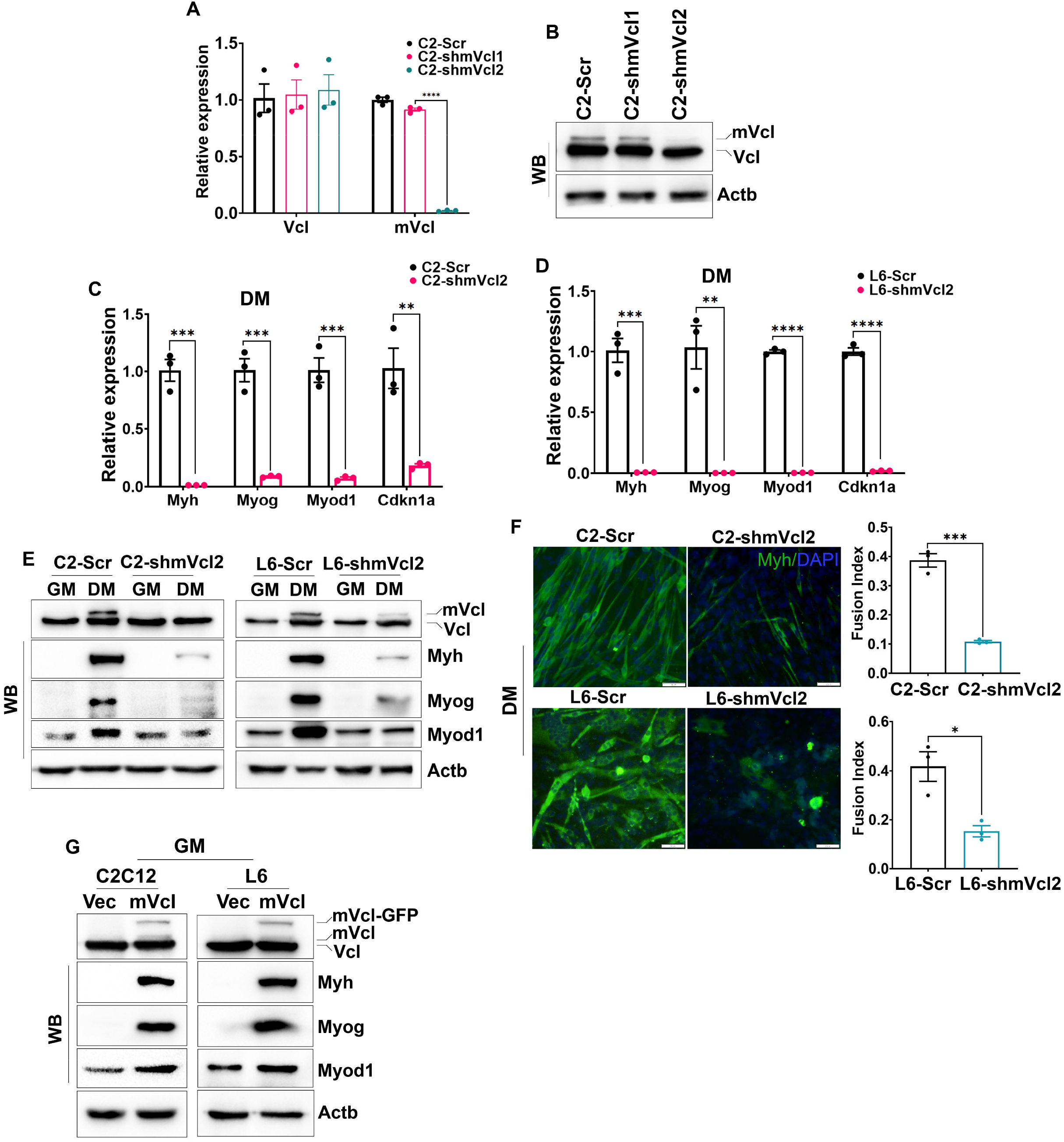
Metavinculin depletion shows defective muscle differentiation, See also in Figure S3. (A and B) Metavinculin depletion in C2-shmVcl1, C2-shmVcl2 and C2-Scr cells that stably express shRNAs targeting metavinculin or scramble is shown in RT-qPCR (A) and western blot (B) analysis. (C and D) Myogenic genes transcripts are reduced under DM conditions in metavinculin depleted C2-shmVcl2 and L6-shmVcl2 cells compared to control C2-Scr or L6-Scr is shown in a bar graph. (E) Western blot showing metavinculin depletion inhibits myogenic proteins induction in C2-shmVcl2 and L6-shmVcl2 cells. (F) Representative images of Myh immunostaining (green) and DAPI counter stained nuclei (blue) in C2-shmVcl2, C2-Scr, L6-shmVcl2, L6-Scr cells in DM conditions and bar graph showing fusion index. (G) Western blot showing Myod1, Myog (early) and Myh (late) myogenic markers under growth promoting conditions in C2C12 and L6 cells having ectopic metavinculin or vector. Actb is a loading control in western blot images. Data are shown as mean value ±SEM, n=3. p value * ≤0.05; **≤0.01; ***≤0.001; ****≤0.0001.

Further, we have analyzed the whole transcriptomic changes by metavinculin depletion using RNA-Seq analysis. We have isolated total RNA from C2-shmVcl2 and C2-Scr cells cultured under differentiation conditions using the Trizol method in biological triplicates. The libraries are synthesized from these RNAs and paired-end sequencing is performed. Here, we have found 1,387 differentially expressed genes (DEGs) (FDR <0.05) includes 809 upregulated and 578 downregulated genes in metavinculin-depleted C2-shmVcl2 compared to control C2-Scr cells using edgeR (Figure 4A and 4B). Randomly picked upregulated (Klhl13, Heatr9, Cxcl10, Ccl5, Acod1, Slurp1) and downregulated (Ccn2, Tagln, Actg2, Myl9, Car3) DEGs are further validated and analyzed for their differentiation-specific expression changes by RT-qPCR analysis. Here, we have found decreased level of Klhl13, Ccl5, Acod1, Slurp1 and increased level of Heatr9, Cxcl10 during skeletal muscle differentiation and they are further increased by metavinculin depletion in C2-shmVcl2 cells (Figure 4C). In other hand downregulated DEGs Ccn2, Tagln, Actg2, Myl9, Car3 expressions are upregulated during normal skeletal muscle differentiation and metavinculin depletion inhibits the same (Figure 4D). Gene ontology (GO) of the differentially expressed genes was analyzed into various functional categories including Biological Process (BP) and Cellular Components (CC) using the online tool ShinyGO 0.80. Upregulated genes are mostly associated with the various stress-responsive immune modulatory GO terms under the BP category (Figure 4E). Further, the CC category of upregulated genes showed the association of DNA replication fork, preinitiation, MCM, and CMG complexes, which suggest DNA replication during the cell cycle (Figure S4A). Similar results were also observed in the KEGG eukaryotic DNA replication complex (Figure S4B). In addition, G1/S-specific cyclin Ccnd1 and G2/M-specific cyclin Ccnb1 are also appeared in upregulated DEGs and further analysis of FPKM data revealed more than two-fold upregulation of Ccna2, Ccnb2, Ccne2 and Ccnf cyclins (Figure S4C, Table S3). These findings further suggest that induction of cell cycle in metavinculin-depleted cells under differentiation-promoting conditions. Besides, BP analysis of downregulated genes showed that all the top ten statistically significant GO terms enriched with muscle contraction, differentiation, and development, in particular striated muscle (Figure 4E). Actin filament-based process and regulation, actin cytoskeleton regulation are also associated with the downregulated DEGs (Figure 4E). Additionally, tube morphogenesis and development was also significant GO terms associate with downregulated genes (Figure 4E). Further, the downregulated DEGs are associated with typical muscle components like myofilament, actin filament bundle, actomyosin, myofibril, I band, and Z disc (Figure S4A). Focal adhesion, adherence junction, cell-substrate junction, and anchoring junction are notable GO terms associated with downregulated genes (Figure S4A). Hence, GO analysis of downregulated DEGs showed that metavinculin depletion severely impedes the genes associated with muscle differentiation and functions in addition to cytoskeletal network genes. In a parallel KEGG analysis, downregulated DEGs are associated with dilated, hypertrophic cardiomyopathies and Adrenergic, Wnt and MAPK signaling (Figure S4D). Interestingly, KEGG Wnt-signaling pathway enriched DEGs are mostly involved in β-catenin independent non-canonical Planar Cell Polarity (PCP) and Wnt/Ca^2+^ pathways (Figure S5A). Further, we have also observed a significant reduction of FPKM of non-canonical Wnt7b in C2-shmVcl2 cells compared to C2-Scr cells after removing the outlier and another non-canonical Wnt11 also showed more than 2-fold reduction of FPKM however it is not statistically significant (Figure S5B). Interestingly, Wnt10a, a canonical Wnt-pathway ligand appeared in upregulated DEGs (Table S3). Moreover, these observations indicate that metavinculin-depletion promotes canonical and inhibits non-canonical Wnt-signaling under differentiation conditions (Figure S5, Table S3). Overall, the transcriptome changes in C2-shmVcl2 cells suggest that metavinculin depletion fails to induce growth arrest and subsequent skeletal muscle differentiation perhaps due to disrupted Wnt-pathways (Figure 4).

**Figure 4:**
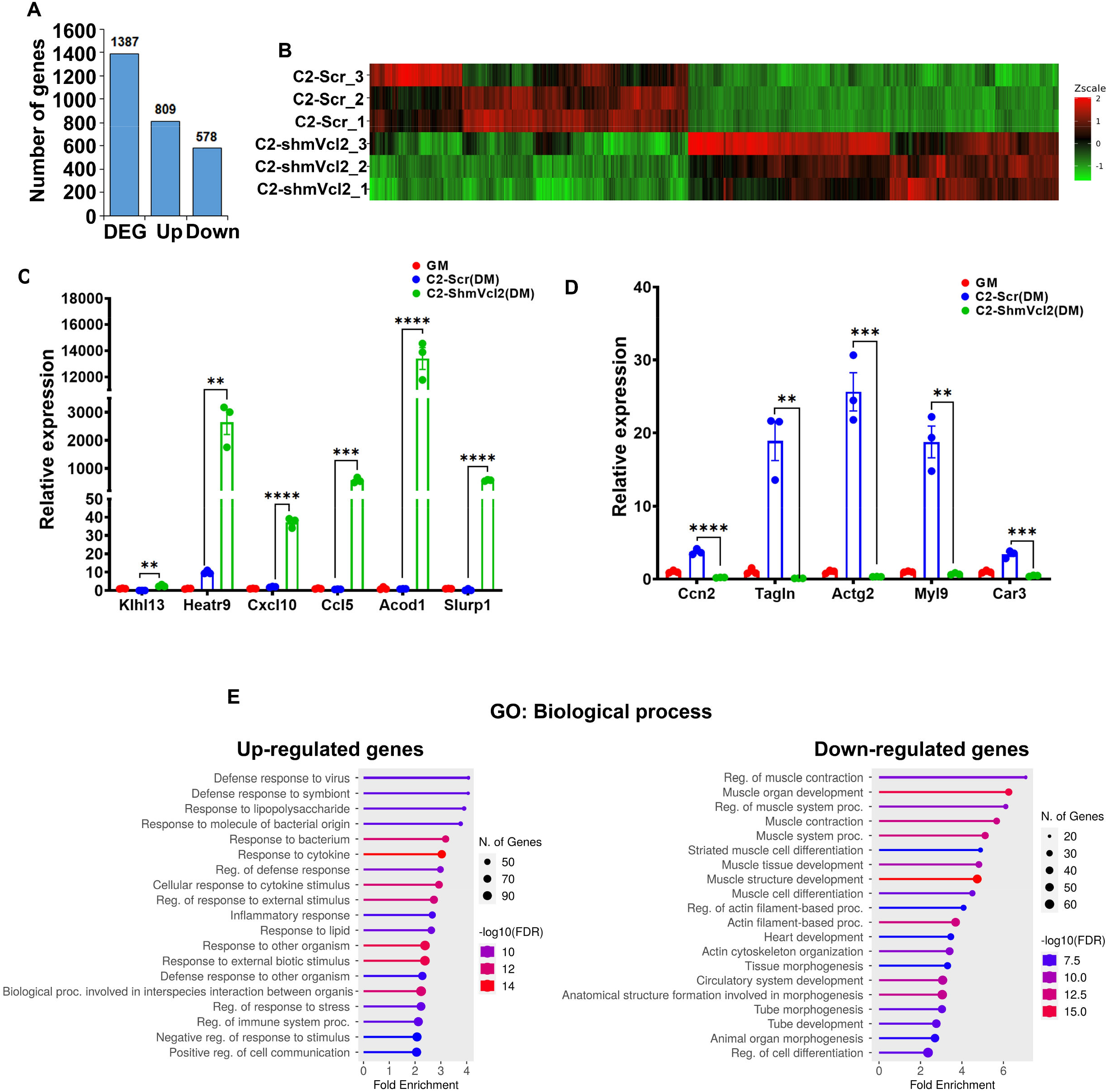
Altered transcriptomic landscape in metavinculin depleted cell, See also in Figures S4-S5, Table S3. (A and B) Differentially expressed genes between metavinculin depleted C2-shmVcl2 and scramble control C2-Scr cells shown in a bar graph (A) and a color scale in the heat map showing normalized expression values (B). (C and D) Bar graph showing differentiation-specific expression and validation of randomly picked up (C) and down (D) regulated DEGs by RT-qPCR analysis. (E) Bar plot showing Gene ontology (GO): Biological Process terms significantly associated with up and down regulated DEGs. Data are shown as mean value ±SEM, n=3. p value **≤0.01; ***≤0.001; ****≤0.0001.

### Canonical and non-canonical Wnt-signaling differentially regulates skeletal muscle differentiation

Studies showed the importance of Wnts in muscle development however most of them are focused on the muscle progenitor maintenance with limited evidence in skeletal muscle differentiation.^72–81^ Here, metavinculin depletion severely disrupts skeletal muscle differentiation with an altered Wnt-signaling pathway (Figures 3, 4 and S5, Table S3). To confirm the significance of Wnt-activation in muscle differentiation, first we have analyzed the expression of Wnt-target genes in growth and differentiation conditions. RT-qPCR results revealed that canonical Wnt-targets Myc, Ccnd1, Ascl2, Id2, Tcf4 and Mmp9 are reduced in cells cultured under DM conditions compared to GM (Figure 5A). Then we have analysed β-catenin (Ctnnb) expression in C2-shmVcl2 and C2-Scr cells under various culturing conditions. In scramble control cells, β-catenin expression present in GM condition is diminished in DM condition whereas β-catenin level is retained in metavinculin-depleted cells (Figure 5B). Since vinculin directly binds with β-catenin to stabilize E-cadherin at the epithelial cell surface^82^ we have verified the similar vinculin/β-catenin interaction during skeletal muscle differentiation by immunoprecipitation analysis. The results revealed vinculin direct interaction with β-catenin in growth conditions and it is absent during differentiation probably due to decreased β-catenin level (Figure 5B and 5C). This interaction is retained in metavinculin-depleted C2-shmVcl2 cells that showed differentiation defect (Figure 3 and 5C). To further explore canonical Wnt-signaling mediated differentiation inhibition, we have treated the cells with the reported canonical Wnt-activator BML-284^83^ and analyzed its effect on differentiation potential. BML-284 treatment activated the canonical Wnt-pathway target genes Myc, Ccnd1, Id2, Tcf4, and Mmp9 in C2C12 cells (Figure 5D). Further, BML-284 treatment completely inhibited the induction of growth arrest gene Cdkn1a and myogenic genes Myh, Myog, Myod1, in C2C12 cells during differentiation (Figure 5E). The levels of Myod1, Myog, and Myh proteins are also inhibited by BML-284 treatment (Figure 5F). Wnt-activation also reduced Myh positive cells and myotube formation in C2C12 cells under DM conditions compared to vehicle-treated cells (Figure 5G). Since canonical Wnt-activation negatively regulates skeletal muscle differentiation the observed differentiation defect in metavinculin depleted cells could be due to sustained canonical Wnt-signaling by β-catenin stabilisation (Figure 5).

**Figure 5:**
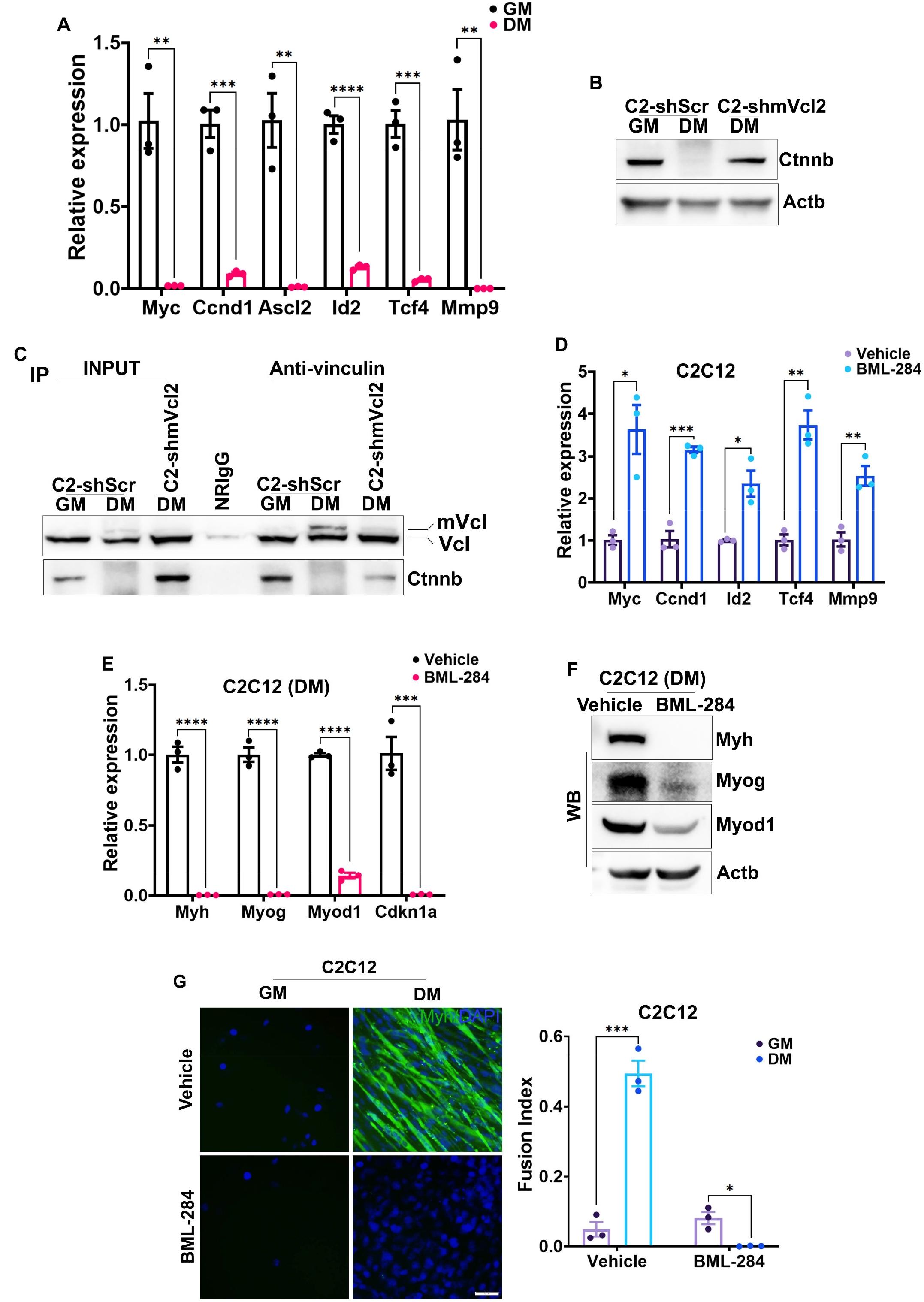
Canonical Wnt-activation negatively affects skeletal muscle differentiation. (A) Relative expression of canonical Wnt target genes Myc, Ccnd1, Ascl2, Id2, Tcf4 and Mmp9 in C2C12 cells cultured under various conditions. (B) Western blot showing canonical Wnt mediator Ctnnb (β-catenin) protein level in C2-Scr and C2-shmVcl2 cells cultured as indicated. (C) Co-immunoprecipitation analysis showing direct vinculin and Ctnnb interaction in C2-Scr and C2-shmVcl2 cells grown under indicated conditions. (D) Bar graph showing RT-qPCR validation of Wnt target genes activation by canonical Wnt activator BML-284 in C2C12 cells. (E) Relative expression of myogenic differentiation associated genes in C2C12 cells treated with BML-284 or vehicle in DM media shown in RT-qPCR. (F) Myogenic protein expression in BML-284 or vehicle treated C2C12 cells cultured under DM conditions is shown in western blot. (G) Representative images of Myh stain (green) and DAPI (blue) in C2C12 cells treated with BML-284 or vehicle. Fusion index is shown in a bar graph. Actb act as a loading control in western blot. Data are shown as mean value ±SEM, n=3. p value * ≤0.05; **≤0.01; ***≤0.001; ****≤0.0001.

Wnt-activation inhibits the Cdkn1a expression which is necessary to induce growth arrest during skeletal muscle differentiation (Figure 5E).^12,69^ As Wnt-activation prevents skeletal muscle differentiation its inhibition may induce differentiation (Figure 5E-5G). To confirm this, we have inhibited the canonical Wnt-pathway using a potent Wnt-inhibitor KY02111^84^ in C2C12 cells (Figure 6A). KY02111-treatment induced Cdkn1a transcripts along with Myod1, Myog, and Myh under the growth promoting conditions compared to vehicle treatment (Figure 6B). We have also confirmed the induction of myogenic genes at the protein level (Figure 6C). KY02111-treatment increased Myh-positive C2C12 cells at growth condition, however, they did not show any fusion (Figure 6D). Since canonical Wnt-inhibition is essential for skeletal muscle differentiation (Figure 6A-6D) it may recover differentiation defect in metavinculin-depleted cells. As expected, KY02111-treatment upregulates Cdkn1a, Myod1, Myog, and Myh mRNAs expression in C2-shmVcl2 cells compared to vehicle treated cell (Figure 6E). Myh, Myog, and Myod1 levels are also induced in these cells by KY02111 (Figure 6F). Increased myotube formation and fusion index in metavinculin-depleted cells are also noted in KY02111 treated cells (Figure 6G). Thus, these results revealed that canonical Wnt-inhibition can induces skeletal muscle differentiation and rescue differentiation in metavinculin depleted cells (Figure 6).

**Figure 6:**
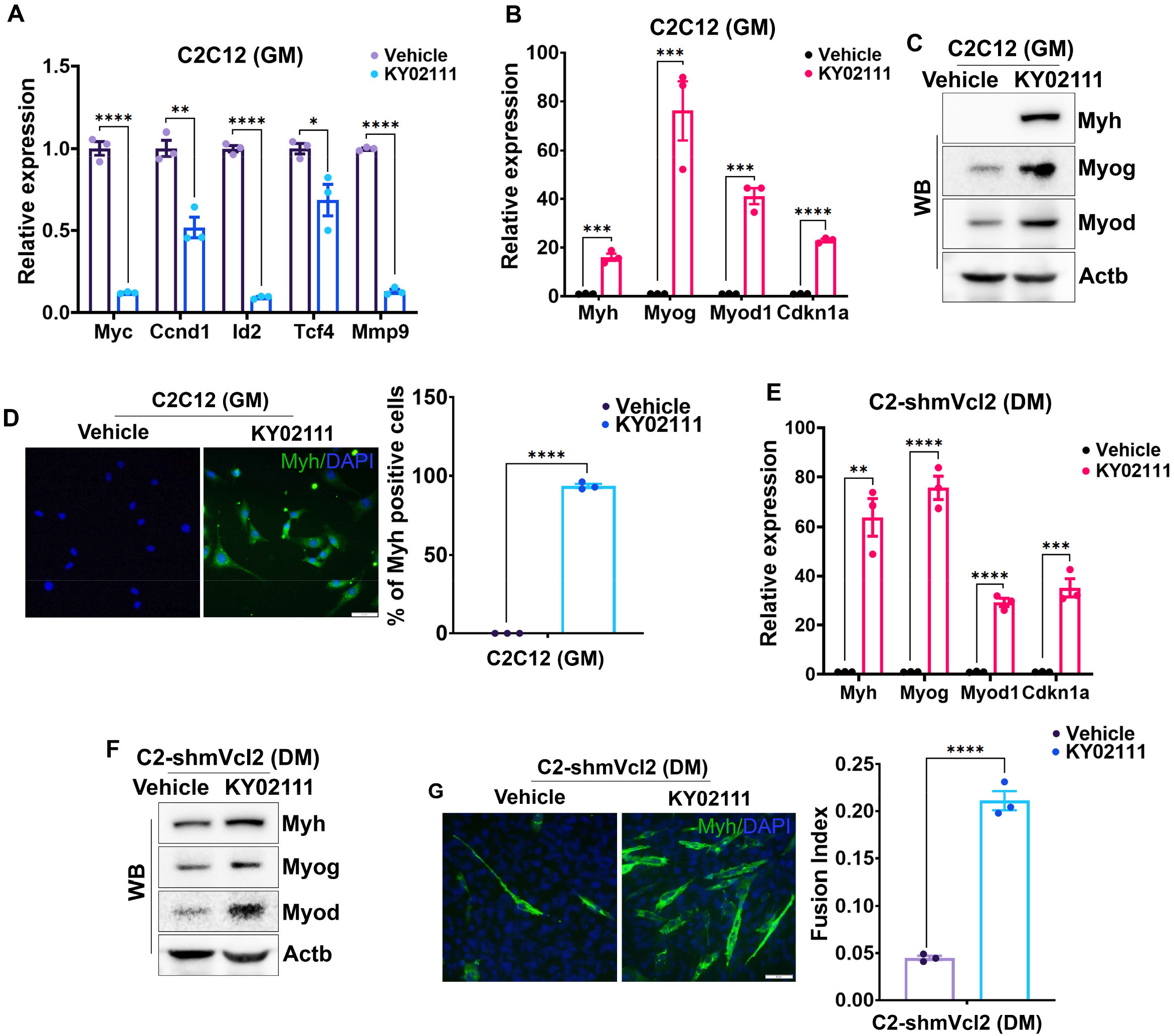
Canonical Wnt-inhibition induces skeletal muscle differentiation program in metavinculin depleted cells. (A) RT-qPCR showing the expression of Wnt target genes in C2C12 cells treated with KY02111. (B) Relative expression of skeletal muscle differentiation marker genes in C2C12 cells treated with KY02111 in growth condition is shown in a bar graph. (C) Western blot showing the induction of myogenic proteins by KY02111 treatment in C2C12 under growth conditions. (D) Representative immunofluorescence images showing Myh positive cells (green) and nuclei (blue) in KY02111-treated C2C12 and its percentage is shown in a bar graph. (E) RT-qPCR analysis of myogenic genes in metavinculin depleted C2-shmVcl2 cells treated with KY02111 in DM conditions. (F) Western blot showing muscle specific genes in KY02111-treated C2-shmVcl2 cells under DM. (G) Representative images of Myh staining (green) and nuclear stain DAPI (blue) in metavinculin-depleted C2C12 cells treated with KY02111 in DM. Fusion index is shown as bar graph. Actb act as a loading control in western blot. Data are shown as mean value ± SEM, n=3. p value * ≤0.05; **≤0.01; ***≤0.001; ****≤0.0001.

Metavinculin-depletion inhibits the non-canonical Wnt-pathway while activating the canonical pathway (Figure S5A, Table S3). Inhibition of non-canonical Wnt7b in metavinculin-depleted cells (Figure S5B) is further confirmed by RT-qPCR (Figure 7A). Antagonism between non-cantonical and canonical Wnt-signaling is reported in several studies and we have also found induction of non-canonical Wnt7b during canonical Wnt-inhibition in C2C12 cells (Figure 7B).^85–87^ Further, we have observed induction of Wnt7b protein in C2C12 and L6 cells during differentiation (Figure 7C). Differentiation-specific Wnt7b induction is inhibited in metavinculin-depleted C2-shmVcl2 and L6-shmVcl2 cells (Figure 7A and 7C). Since canonical Wnt-inhibition can activate non-canonical Wnt-signaling we have analyzed the status of non-canonical Wnt7b in KY02111-treated metavinculin-depleted cells (Figure 7B).^85–89^ KY02111-treatment induced Wnt7b mRNA and protein levels in C2-shmVcl2 compared to vehicle while restoring skeletal muscle differentiation (Figure 6E-6G, 7D and 7E). Similarly, ectopic Wnt7b expression also likely to restore skeletal muscle differentiation these cells (Figure 7F and 7G). As expected, retroviral-mediated ectopic Wnt7b expression restored of myogenic genes Myh, Myog, Myod1 and Cdnk1a in C2-shmVcl2 cells, however, the induction of late myogenic marker Myh is less compared to early myogenic genes (Figure 7H). Further, the restoration of myogenic genes are also reflected in their protein levels (Figure 7I). Ectopic Wnt7b expression is also improved myotube formation and fusion index in metavinculin-depleted cells compared to vector control (Figure 7J). Collectively, these data revealed that canonical Wnt-inhibition or non-canonical activation can restore skeletal muscle differentiation in metavinculin depleted cells (Figure 6 and 7).

**Figure 7:**
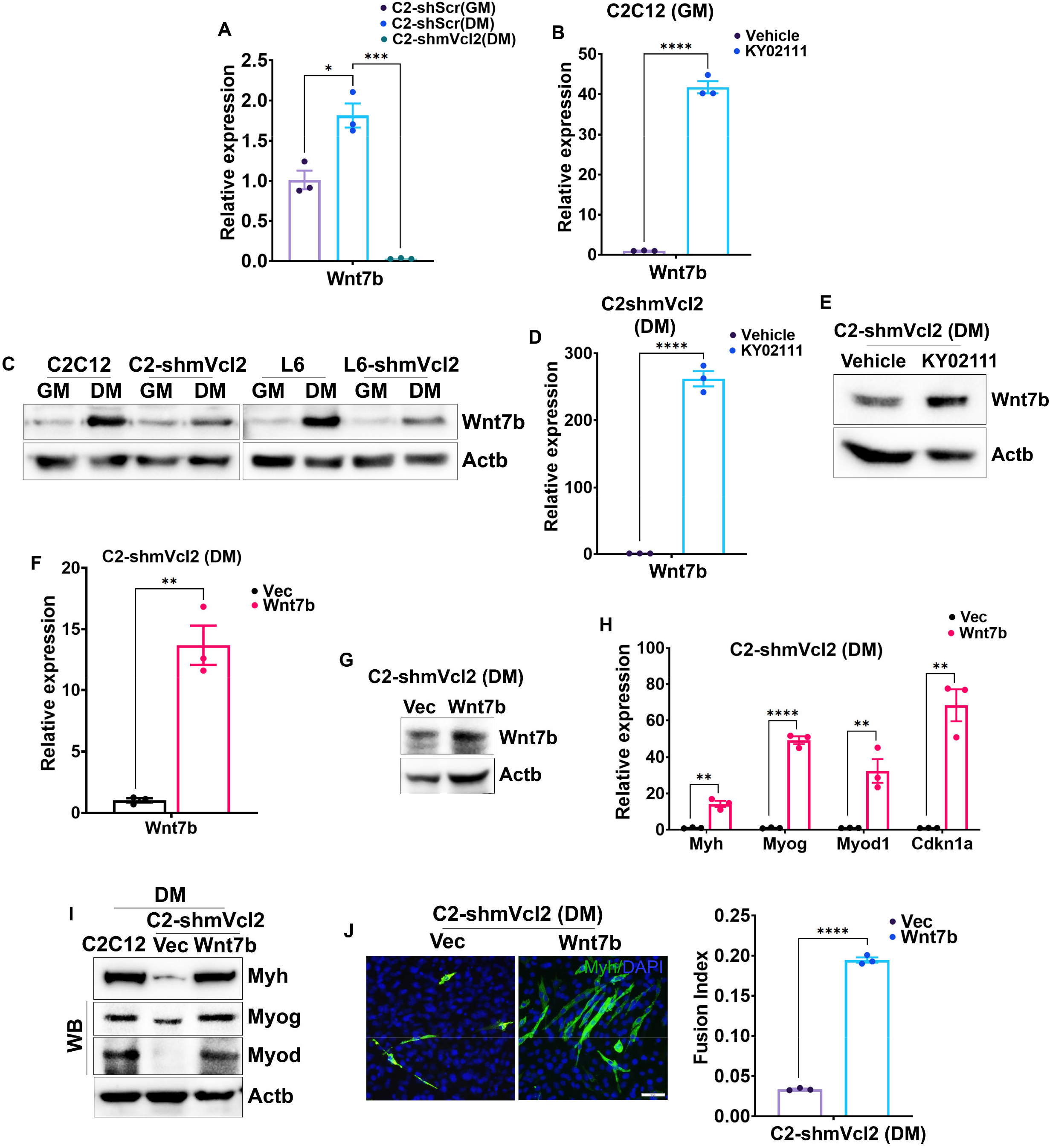
Ectopic expression of non-canonical Wnt7b enhances skeletal muscle differentiation in metavinculin depleted cells. (A) Bar graph showing relative expression of Wnt7b in C2-Scr and C2-shmVcl2 cells cultured under various conditions. (B) Relative expression of Wnt7b in KY02111-treated C2C12 cells is shown in a bar graph. (C) Induced Wnt7b protein during C2C12 and L6 cells differentiation is inhibited by metavinculin depletion as shown in western blot. (D and E) KY02111-treatment induced Wnt7b expression at RNA (D) and protein (E) level in C2C12 cells is shown by RT-qPCR and western blot respectively. (F and G) Ectopic expression of Wnt7b is verified in RT-qPCR (F) and western blot (G) analysis. (H) Bar graph showing increased the expression of myogenic marker genes in C2-shmVcl2 cells by ectopic Wnt7b expression. (I) Western blot showing upregulation of myogenic proteins by Wnt7b ectopic expression in C2-shmVcl2 cells compared to vector control. (J) Representative pictures of Myh stain with DAPI counter stain in C2-shmVcl2 cells that has ectopic Wnt7b or vector. Fusion index in these cells are represented in a bar graph. Actb act as a loading control in western blot. Data are shown as mean value ± SEM, n=3. p value * ≤0.05; **≤0.01; ***≤0.001; ****≤0.0001.

## DISCUSSION

### Metavinculin is differentially expressed during myoblast proliferation and differentiation

Metavinculin express along with vinculin at sub-stoichiometric in skeletal muscle as well as smooth, cardiac muscles (Figure 1A).^37–39,45,47,68^ However, proliferation dependent reduction of metavinculin level reported in primary cells isolated from human aorta and chicken gizzards and completely absent when they sub-cultured.^37,43,45^ Similarly, we have found metavinculin expression only in differentiated myotubes not in the proliferating myoblasts while vinculin level remains unaltered (Figure 1B-1D). Proliferating myoblasts first exit from the cell cycle and undergoes terminal muscle differentiation by activation of various myogenic genes.^12,69^ Differentiation signal leads to master myogenic regulator Myod1 activation which in turn activates its targets Cdkn1a (a growth arrest gene) followed by Myog by altering their epigenome.^5,8–10^ Later, Myod1 and Myog collaborate to activate terminal myogenic genes.^3,11^We have found metavinculin isoform expression gradually increased during in vitro skeletal muscle differentiation with no change in vinculin level (Figure 1D). After muscle injury, activated resident satellite cells extensively proliferate and undergo differentiation program to complete muscle regeneration process.^90,91^ Here, we have found a direct correlation of metavinculin expression with skeletal muscle differentiation during regeneration in the mouse cryoinjury model (Figure 1E-1G); however, its expression is transiently reduced at the early stages of regeneration where excessive cell division takes place (Figure 1G). Hence, it appears that metavinculin isoform switching is controlled based on the status of muscle cell proliferation or differentiation (Figure 1B-1G).^37,39,45^

### Rbfox1 regulates metavinculin splicing during skeletal muscle differentiation

Alternative splicing is also major contributor for the diverse muscle proteome in addition to myogenic transcription factors.^1,6,14,16,92^ Alternative transcript usage gives spatiotemporal expression of unique isoforms during skeletal muscle differentiation and it strongly correlates with the various splicing regulators, including Rbfox, Celf, Mbnl and Ptb expression.^64–67^ Among these factors, Rbfox family may collaborate with other splicing regulators to establish muscle-specific splicing events during muscle differentiation as they exhibit high tissue specificity (Table S1).^19,71^ Rbfox1 and Rbfox2 are the major Rbfox family members expressed in muscle cells. Rbfox1 expression is induced during myotube formation while Rbfox2 expression remains unaltered and the same is also reported by others (Figure 2A and 2B).^19,71^ We did not find consistent change in Rbfox3 level during muscle differentiation as it is involved in neuronal development (Figure 2A and 2B). Rbfox family include or exclude exons depending on the location of its cis-regulatory elements, in specific, upstream location excludes and the downstream location includes the associated cassette exon.^93^ Metavinculin is a larger vinculin isoform specifically expressed in differentiated muscle cells and its expression is reduced in proliferating cells which coincides with Rbfox1 expression (Figure 1A-1C, 2A and 2B).^37,39,45^ Although both Rbfox1 and Rbfox2 can bind in vitro to metavinculin flanking introns the binding is enriched at downstream intron in myotubes (Figure 2C and 2D). Moreover, Rbfox1 regulates metavinculin expression over Rbfox2 as shown by depletion experiments (Figure 2E-2G) which is strengthened by a recent report.^94^ Further, crosslink-RIP analysis confirmed the differentiation-specific switching of Rbfox1 from 5’-to 3’-metavinculin flanking intron (Figure 2H-2J). While ectopic Rbfox1 expression showed induction of metavinculin transcripts under growth-promoting conditions however it is not reflected at protein level which suggested the unstable metavinculin protein in proliferating cells (Figure 1B, 1D and 2K-2N).^37,43,45^

It was reported that Rbfox1 or Rbfox2 depletion does not affect early stages of muscle differentiation and it affected the myotube formation.^19,71^ In contrast, we have observed Rbfox1-depletion severely inhibited the expression of early myogenic genes Myod1, Myog in addition to terminal differentiation marker Myh whereas certain level of these myogenic genes are retained in Rbfox2-depleted cells and this difference observation is probably due to transient or stable nature of depletion (Figure S2C and S2D).^19,71^ Although both Rbfoxs are necessary for skeletal muscle differentiation Rbfox1 contributes more during differentiation as its ectopic expression can induce differentiation associated genes even at growth promoting conditions (Figure S2E, S2F, 2O and 2P). Interestingly, we have found Rbfox1-depletion negatively and Rbfox2-depletion positively correlated with each other’s expression (Figure 2E). Thus, it appears that there is a negative feedback inhibition between Rbfox2 and Rbfox1 expression during skeletal muscle differentiation and it will be interesting to explore further. Nevertheless, these results suggest that induced Rbfox1 expression critically regulates metavinculin alternative splicing during differentiation by preferentially binding to its downstream flanking intron (Figure 2).

### Metavinculin critically regulates muscle cell proliferation and differentiation

Vinculin links the integrin adhesion complex to the filamentous actin network and evidences suggest that metavinculin play a similar role along with vinculin with enhanced force transduction, however, its muscle-specific remains to be identified.^34,42,44,58,61,63,95^ Here, downregulated DEGs in metavinculin-depleted cells are associated with focal adhesion, adherens, anchoring and cell-substrate junctions in skeletal muscle cells as metavinculin is necessary to stabilize these complexes similar to vinculin (Figure S4A). The major functional difference reported between vinculin and metavinculin lays in the actin bundling.^42,58,61–63^ Metavinculin may fine-tune actin organisation in skeletal muscles as downregulated DEGs in metavinculin-depleted cells associates with actin bundling terms (Figure S4A).^58,59,61,63^

Moreover, we have identified the unique role metavinculin in skeletal muscle differentiation through depletion and ectopic expression studies. Specifically, metavinculin depletion inhibited early and late genes involved in muscle differentiation (Figure 3A-3F). Interestingly, ectopic expression of metavinculin is able to induce these genes even in growth-promoting conditions (Figure 3G). Transcriptome analysis in metavinculin-depleted cells also confirmed the downregulation of genes associated with muscle development and contraction (Figure 4C). In particular, downregulated DEGs in metavinculin-depleted cells associate with cellular compounds actomyosin, myofibril, contractile fiber, sarcomere, I band, and Z disc (Figure S4A). Vinculin knockout is embryonic lethal due to multiple defects in the heart, neural tube, somites, and limb.^36^ In agreement with this metavinculin depletion also affected the several genes involved in muscle and neural tube development (Figure 4E). Mutations at the metavinculin-specific exon or loss of expression reported to be involved in various cardiomyopathies.^44,48–53,95,96^ Downregulated DEGs by metavinculin depletion is also associate with these cardiomyopathies (Figure S4D). Surprisingly, the heart muscle sections from muscle-specific metavinculin knockout mice do not show any defect in cardiac tissue architecture, and the expression/localization of the costamere, intercalated disc, and gap junction proteins also remains unaltered although they report metavinculin bears higher molecular forces.^95^ This discrepancy in their observation is because they targeted the upstream region which will not disrupt metavinculin-specific exon 19, as shown in their supplementary figure 10.^95^ However, the evidence from our study supports the metavinculin play a crucial role in establishing myogenic gene expression during differentiation and muscle function (Figure 3 and 4).

Metavinculin isoform switching occurs during muscle differentiation and is absent in proliferating cells (Figure 1B and 1C).^38,43,45^ Moreover, metavinculin depletion inhibits the expression of growth arrest gene Cdkn1a which is necessary to induce cell cycle exit during skeletal muscle differentiation (Figure 3A-3D).^12,69^ Further, metavinculin depletion upregulated the genes involved in various stress-responsive pathways, DNA replication complexes, and several cyclins (Figure 4E, S5). Thus, it appears that metavinculin expression is needed for growth arrest prior to skeletal muscle differentiation as its absence retains the cells in a proliferative stage even under differentiation-promoting conditions. Taken together, these results indicate that metavinculin is critical for muscle cells proliferation and differentiation decisions.

### Metavinculin switches canonical to non-canonical Wnt-signaling during muscle differentiation

Wnt-signaling plays a crucial role in stem cell proliferation, differentiation, and self-renewal during embryonic development as knockout of Wnt-pathway genes showed embryonic lethality with multiple tissue defects including muscle.^72,73,75,77,79,80^ In the current study, metavinculin-depletion inhibited the genes associated with muscle development along with Wnt-pathway deregulation (Figure 3, 4F, S4 and S5, Table S3). Several Wnt ligands controls MRFs expression during muscle progenitor cell proliferation and differentiation.^72,77–79^ Specifically, canonical Wnt1, 3a, 4, and 6 maintain the expression of Pax3 and Pax7 in muscle progenitors.^97^ Wnt1 and Wnt3a double knockout mouse embryos failed to form proper dermomyotome due to reduced expression of Myf5.^77^ Further, Wnt1 regulates Myf5 expression and non-canonical Wnt7a activates Myod1 in epaxial and hypaxial muscle respectively.^72,74^ Canonical Wnt mediator β-catenin inhibition is required for proper skeletal muscle differentiation as its activation by BML-284 prevents muscle differentiation (Figure 5). We have found direct binding of β-catenin with vinculin during myoblast proliferation and it diminished during differentiation (Figure 5C). Restored β-catenin level by vinculin/β-catenin interaction may responsible for the differentiation defects observed in metavinculin-depleted cells (Figure 3, 4F, 5B and 5C). Further, canonical Wnt inhibition by KY02111 induce myogenic genes even in growth-promoting conditions and restores myogenic differentiation in metavinculin-depleted cells (Figure 6). Hence, these evidences suggest that metavinculin destabilizes β-catenin by interfering β-catenin/vinculin association during skeletal muscle differentiation.

Non-canonical Wnt5a, Wnt5b, Wnt7a, Wnt7b are important for various stages of muscle regeneration.^73,98,99^ Moreover, Wnt11-mediated Wnt/PCP pathway is necessary for embryonic muscle fibre formation.^76^ A recent study demonstrates that Wnt7b expression in muscle stem cells is repressed by the cis-acting long noncoding RNA Lnc-Rewind and it is relived during differentiation.^100^ We have also found the induction of non-canonical Wnt7b during skeletal muscle differentiation and metavinculin-depletion inhibited the same (Figure 7A and 7C). KY02111 treatment activates non-canonical Wnt7b and restored myogenic differentiation in metavinculin-depleted cells (Figure 6E-6G, 7B, 7D and 7E). Similarly, Wnt7b induction is observed during myogenic differentiation in canonical Wnt10b^-/-^ myoblasts.^98^ Non-canonical Wnt ligands can alter canonical Wnt-signaling in a spacio-temporal manner.^73,98^ Specifically, non-canonical WNT/Ca2^+^ pathway-mediated activation of NLK and HIPK2 inhibits the transcriptional activity of endpoint Wnt-signaling mediator TCF/LEF by phosphorylation.^86^ In addition, non-canonical Wnt5a is known to inhibit Wnt3a-mediated canonical signaling during hematopoietic stem cell differentiation.^85^ Here, ectopic Wnt7b is sufficient to improve myogenic induction and the fusion index in metavinculin-depleted cells (Figure 7F-7J). In contrast to these observations involvement of canonical Wnt-signalling in inducing muscle differentiation is reported in various studies.^73,78,79,99,101,102^ However, these studies further insist its role in muscle progenitor maintenance not terminal differentiation. Hence, it appears that canonical Wnt-signaling regulates muscle progenitor proliferation and maintenance whereas non-canonical Wnt signals involve in terminal muscle differentiation. To conclude, vinculin/β-catenin association sustains canonical Wnt-signaling in proliferating myoblasts. During skeletal muscle differentiation, upregulated Rbfox1 generates metavinculin isoform which in turn switches canonical to non-canonical Wnt-signaling in establishing skeletal muscle differentiation program.

## LIMITATION OF THE STUDY

Here we have focussed on metavinculin role in skeletal muscle differentiation although we aware that metavinculin expression found in cardiac and smooth muscle cells. The available studies on metavinculin mainly focussed in cardiac muscle function as its mutations reported in cardiomyopathies and other muscle cells left unexplored. This prompts us to explore the role of metavinculin in skeletal muscle differentiation though it may involve in the differentiation of other muscle cells. Muscle-specific or general metavinculin knockouts could have given a detailed information on its tissue specific role which is another limitations of this study. Furthermore, regulation of Wnt-signaling by metavinculin is focussed in this study, however, our transcriptomic studies revealed other pathways like MAPK and Adrenergic signaling may also be regulated by metavinculin. However, these limitations emphasize future research in these directions.

## ACKNOWLEDGEMENTS

This work is supported by grants from Science and Engineering Research Board (SERB), India, file number EEQ/2017/000322, ECR/2016/000265 and University Grants Commission, India UGC/FRP F.NO 4-5 (95)/2014 (BSR)(FRP) to MJ. The authors thank Dr. Ashok Arasu, a former graduate student, for his assistance in cloning the metavinculin shRNAs. The authors greatly acknowledge Dr. Asoke Mal, Dr. Sam J Mathew, Dr. Davide Gabellini for providing crucial reagents. We thanks Central Animal Research Facility, NIMHANS. AM is a Senior Research fellow (ICMR-SRF), 45/4/2020-HUM/BMS, from the Indian Council of Medical Research (ICMR). Graphical abstract created with BioRender.com.

## AUTHOR CONTRIBUTIONS

Conceptualization, M.J.; Data curation, A.M., M.J.; Formal Analysis, A.M., M.J.; Investigation, A.M., M.J.; Methodology, A.M., N.B., M.J.; Validation, A.M., N.B.; Writing-original draft, A.M., M.J.; Writing-review & editing A.M., M.J., N.B.; Project administration M.J.; Supervision, M.J.; Funding acquisition, M.J.

## INCLUSION AND DIVERSITY

We support inclusive, diverse, and equitable conduct of research.

---

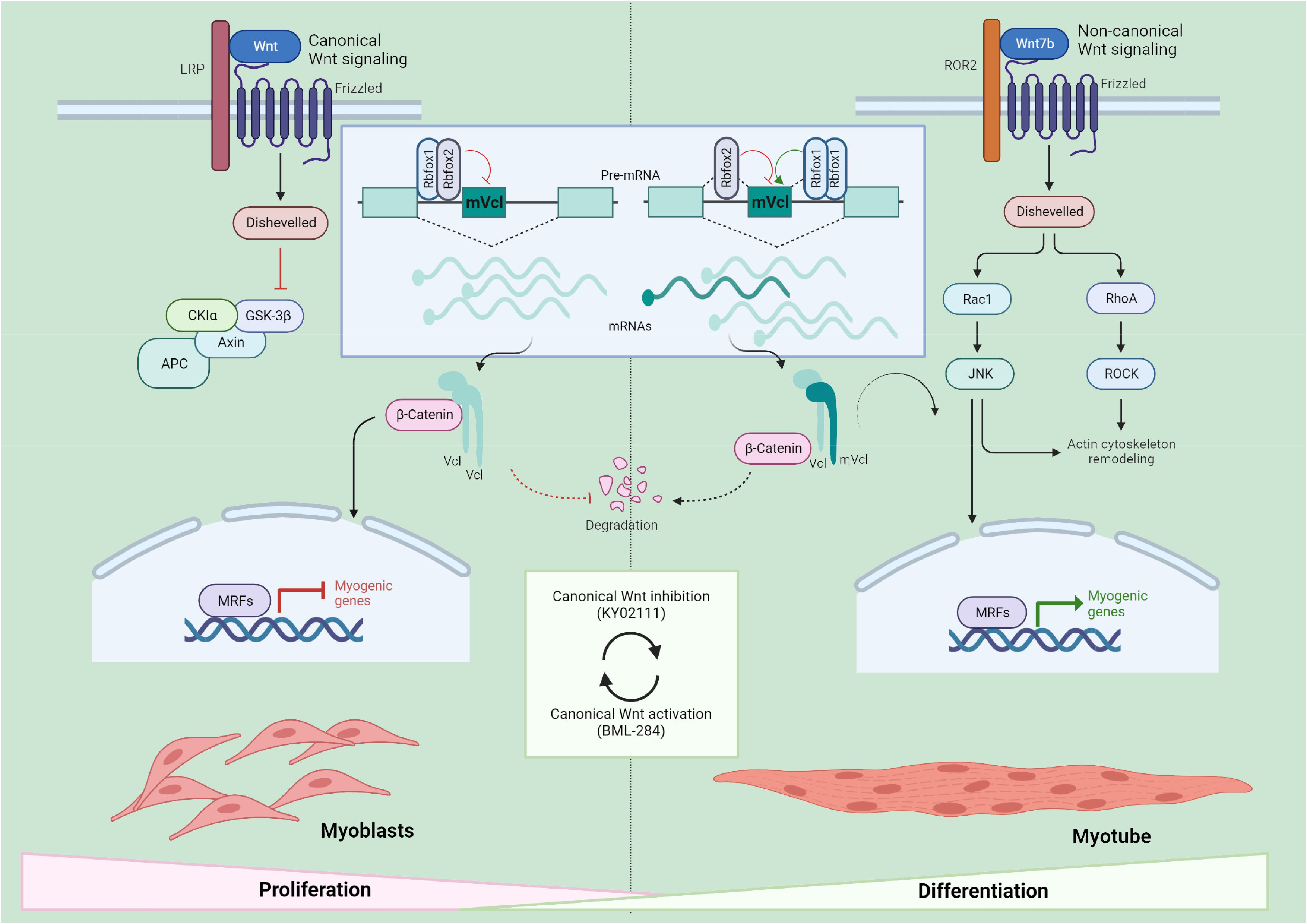

## STAR * METHODS

## RESOURCE AVAILABILITY

### Lead contact

Further information and requests for resources and reagents should be directed to and will be fulfilled by the lead contact, Mathivanan Jothi.

### Materials availability

All unique/stable reagents generated in this study are available from the lead contact with a completed materials transfer agreement.

### Data and code availability

- The RNA-Seq data that support the findings of this study have been deposited on the Gene Expression Omnibus (GEO) and are publicly available. Accession number is listed in the key resources table. All data reported in this paper will be shared by the lead contact upon request
- This paper does not report original code.
- Any additional information required to reanalyze the data reported in this paper is available from the Lead Contact upon request.

## EXPERIMENTAL MODEL AND STUDY PARTICIPANT DETAILS

### Animals

All mouse protocols were in accordance with NIMHANS Institutional Animal Ethics Committee (IAEC) guidelines (Approval number: AEC/74/487/H.G.) which is registered under Committee for Control and Supervision of Experiments on Animals (CCSEA), Ministry of Fisheries, Animal Husbandry and Dairying, Government of India (Registration number: 12/GO/ReBi/S/1999/CPCSEA). Three months old C57BL/6J male mice were used for cryoinjury muscle regeneration model. Cryoinjury was created for 15 seconds in a right tibialis muscle using 5mm sterilized metal rod precooled in dry ice and uninjured contralateral muscle was used as a control. Animal weights were monitored on daily basis, uninjured and regenerating muscle tissues were harvested at 3, 5, 10, 15 and 20 days of post injury after sacrifice for further analysis.

### Cell lines

The C2C12, L6, HSMM myoblasts and 293T cells were used in this study. These cells were cultured in a Dulbecco’s modified Eagle’s medium (DMEM) containing either 20% or 10% Fetal Bovine Serum (FBS) and antibiotic-antimycotic solution (Gibco). Myoblast cultured in DMEM media contains 20% Fetal Bovine Serum (FBS). These cells were induced to differentiate into myotubes by switching the cells into differentiation media containing 2% horse serum (Gibco) for 48 hours (C2C12) or 10days (L6, HSMM). Wnt activator BML-284 (5 mM) or inhibitor KY02111 (10 mM) and vehicle was treated in indicated culturing conditions and refreshed during media change. Stable polyclonal population of C2-shmVcl1, C2-shmVcl2, C2-shRbfox1, C2-shRbfox2, C2-shScr, L6-shScr, L6-shmVcl2 were generated through lentiviral transduction followed by puromycin selection. Similarly C2-Rbfox1 cells were generated using retroviral transduction. All the cells were maintained at 37°C, 5% CO2 in a humidified atmosphere. The cell line authentication is not done in rodents (C2C12, L6) and primary human (HSMM) cell lines.

## METHOD DETAILS

### Molecular cloning

shRNA oligos targeting Rbfox1, Rbfox2, metavinculin and scramble were designed in our laboratory and synthesized from the Integrated DNA Technologies (IDT) along with antisense oligos the sequence details were given in key resources table. The sense and antisense oligos were annealed and cloned into the lentiviral pLKO.1-TRC or pLSL vector. Specifically, Rbfox1, Rbfox2 and Scramble shRNAs were cloned between EcoRI and AgeI sites in pLKO.1-TRC vector. Metavinculin targeted two shRNAs (shmVcl1 and shmVcl2) and scramble were cloned into pLSL into BamH1 and EcoR1 sites. For GST fused Rbfox1 and Rbfox2 recombinant proteins, we have subcloned respective cDNAs fragments digested from pEGFP-Rbfox1 and pEGFP-Rbfox2 with BglII and SalI into pGEX6P-1 plasmid between BamHI and SalI sites. Wnt7b coding sequence was PCR amplified from cDNAs generated from C2C12 cells using specific primers contains BamHI or EcoRI restriction sites and cloned into T vector pMD20. Further, Wnt7b cDNAs was subcloned into pBabe-Puro vector at BamHI and EcoRI sites. Similarly, Rbfox1 cDNA was subcloned from pEGFP-Rbfox1 plasmid. All the inserts in the cloned plasmids and their orientations were confirmed by Sanger’s sequencing.

### Western blot analysis

Twenty milligrams of heart, intestine and tibialis anterior muscle tissues from C57BL/6J mouse and regenerating tissue obtained at various time points were homogenized in a cell lysis buffer containing with 1X protease and phosphatase inhibitors and sonicated to get a tissue extracts. Similarly, total extracts were prepared from cell lines using same lysis buffer. Forty microgram of the obtained tissue or cellular extracts were used for western blot analysis. Briefly, these extracts were separated on SDS-PAGE, transferred to PVDF membrane, incubated with specific primary antibodies and probed with respective secondary antibodies. The primary and secondary antibodies used in a western blot analysis are given in the key resource table. The signal was detected and visualized by ChemiDoc XRS+ (Biorad) or Alliance Q9 (UVITEC) using Immobilon® western HRP substrate (Merck).

### Quantitative RT-PCR

Two microgram of total RNAs isolated from tissues or cells using TRIzol method were converted to cDNA using RevertAid first strand cDNA synthesis Kit (Thermo Scientific) according to the manufacturer instructions. qRT-PCR from these cDNAs were performed using Maxima SYBR green/ROX qPCR master mix (Thermo Scientific) in a QuantStudio™ 6 Flex Real-Time PCR System (Applied Biosystems) using gene specific PCR primers given in Table S1. The results were analyzed by 2^-ΔΔ^ CT method. The gene expressions were normalized with endogenous control Actb.

### Tissue preparation and histopathological analysis

Muscle regeneration in cryoinjured tissues were analyzed by routine standard histological analysis. Concisely, formalin fixed injured and control muscle tissues were processed in an automatic tissue processor (Tissue-Tek® VIP™ Vacuum Infiltration Processor, Sakura Finetek USA, Inc.) and embedded in paraffin blocks. The tissues in paraffin blocks were subjected to sectioning (3-4 μM) by rotary type of microtome (Leica RM2255 Rotary Microtome, Leica Biosystems). The tissue sections were fixed in a glass slide and stained with hematoxylin and eosin (H&E) and Masson’s trichrome stain separately for histological analysis after deparaffinization and rehydration. The histological features were analyzed from the stained images captured from Olympus BX53 microscope.

### Immunofluorescence

Cells cultured in 35-mm dishes at various conditions were fixed with 4% paraformaldehyde, permeabilized with 0.1% Triton X-100 in PBS and blocked with 3 % control serum. After washing, these cells were incubated overnight with anti-Myh antibody (DSHB). Then, primary antibody solution was removed, the cells were washed and incubated with Alexa-Fluor 488 (green) conjugated secondary antibody (Invitrogen) to detect the bound primary antibody. Nuclei were counter stained with DAPI and the images were acquired from fluorescence microscope (Olympus IX73). The fusion index from the fluorescence images were calculated using Myotube Analyzer tool.^103^

### Recombinant protein purification

GST-Rbfox1, GST-Rbfox2 and GST proteins were purified as mentioned earlier.^8^ Briefly, transformed BL21(DE3) cells with respective pGEX6P-1 plasmids were cultured until OD reaches 0.6 and the heterologous protein expression was induced with 0.25 mM IPTG for 3 hr. Then the bacteria were lysed with lysis buffer (50 mM TrisCl pH 7.5, 120 mM NaCl, 1 mM EDTA 0.5 % NP40 and 1 mM PMSF). The clarified supernatant was incubated with GST-agarose beads (Merck) overnight. Then the recombinant proteins were eluted from washed beads with buffer contains 50 mM Tris pH 8.0, 1 mM DTT and 10 mM reduced glutathione. The recombinant proteins were used for GST-pulldown assay after dialysis.

### GST-pulldown assay

Total RNAs isolated form cells grown under GM and DM were treated with Purelink^TM^ DNAse (Invitrogen^TM^). The RNAs then denatured and renatured by heating 2 min at 90°C followed by gradual cooling to room temperature. Two micrograms of total RNA per pull down were pre-cleared with Glutathione-Agarose beads (Merck) in the assay binding buffer (PBS containing 2 mM MgCl2, 0.2 mM ZnCl2, 1mM DTT, 100 U/ml RNAse Inhibitor, 0.1 mg/ml yeast tRNA (Sigma), 0.05% BSA, and 0.2% NP40). 1/10 of this material was used as an input. After pre-clearing the RNAs were incubated with two micrograms of GST-Rbfox1 or GST-Rbfox2 or control GST proteins at room temperature for 1 hour in the assay binding buffer. Twenty microliter Glutathione-Agarose beads was added and incubated overnight at 4°C to pulldown protein/RNA complex. Then, the beads were washed four times in the same assay buffer supplemented with 150 mM NaCl at room temperature. The RNAs from the bead-protein-RNA complex and inputs were recovered by TRIzol method followed by on-column purification, and the eluted RNAs were converted into cDNA and analyzed by qPCR with specific primers given in Table S1.

### Lenti and retroviral transduction

The viral protocols are reviewed and approved by Institutional Biosafety Committee (IBSC), NIMHANS with the approval number NIMHANS/DO/INSC MEETING/2017 S.No.5 dated 27.07.2017) Retro and lenti viruses were produced in 293T packaging cell lines as described previously.^104^ Here, viral plasmids containing cDNA or shRNA along with packaging plasmids were transiently transfected into 293T cell by calcium phosphate co-precipitation method. Forty eight hours of post-transfection the viral supernatants were passed through 0.45 μm syringe and the diluted viruses were used to infect cells in growth media supplement with 8 μg/ml of fresh polybrene (Merck) for 8 hr for three consecutive times. After 48 hr post-transduction the cells were selected with puromycin until the untransduced cells died or used as transient infection. Viral titers were assessed by GFP florescence in 293T cells transduced with serial dilution of GFP viral stocks.

### Crosslink-RNA immunoprecipitation

The cells grown in appropriate conditions were crosslinked with 4% paraformaldehyde and collected in PBS. The nuclei from these cells were extracted using nuclear extraction buffer (1.28M Sucrose, 40mM Tris-Cl pH: 7.5, 20Mm MgCl2, 4% Triton X-100) and the pelleted nuclei were resuspended in RIP buffer (150 mM KCl, 25 mM Tris, pH 7.4, 5 mM EDTA, 0.5 mM DTT, 0.5% NP40, and 100 U/mL RNase inhibitor). The chromatin was sheared by sonication and clarified by centrifugation. For RIP, 4 μg of anti-Rbfox1 antibody (Abclonal) or control NRIgG was incubated overnight at 4 °C with gentile rotation and 2 % of these samples taken as input prior to antibody addition. Then the samples were incubated with protein A-agarose beads (SCBT) for 2 h at 4 °C to pulldown immune complex. The beads were washed three times with RIP buffer and one time with PBS. Then it was divided into two part for protein and RNA analysis. The presence of Rbfox1 protein in the immune complex and input were analyzed by western blot. The RNAs were isolated from the beads and input using TRIzol method followed by on column purification. These RNAs were used in RT-qPCR analysis as mentioned elsewhere. The primer details are given in the Table S1.

### Chromatin immunoprecipitation (ChIP)

We have followed ChIP protocol as mentioned earlier.^105^ Briefly, the cells grown in appropriate conditions were fixed with 37 % formaldehyde and collected by scraping in ice cold PBS with PMSF. The cells were suspended in SDS lysis buffer with protease inhibitor cocktails and sonicated 15 S each at 50 amplitude for 25 times in a probe sonicator (Qsonica). The fragment size was verified from fifty microliter of sonicated samples. The chromatin (100 μg) was precleared with protein A agarose/Salmon sperm DNA beads (Merck) and 2 % taken as input. Then the chromatin was incubated with anti-Myod antibody and NRIgG antibodies overnight at 4 °C. Next the immune complex was recovered by protein A agarose/Salmon sperm DNA beads incubation for 4 hr at RT and the beads were serially washed with low salt, high salt, LiCl and TE buffers. Then the beads were treated with RnaseA, proteinase K. DNA fragments from the beads and input samples were purified using PureLink™ Quick Gel Extraction and PCR Purification combo kit (Thermo Scientific) after decrosslink. Myod target Myog and Myh promoter regions were detected in these samples by qPCR using primers given in Table S1.

### RNA-sequencing

Total RNAs isolated from C2-shScr and C2-shmVcl2 cells were used for RNA-sequencing analysis. RNA integrity was assessed using an Agilent 2200 TapeStation system with an RNA Integrity Number (RIN) value greater than or equal to 7. The libraries were synthesized from the these RNAs using TruSeq Stranded Total RNA LT sample Prep Kit (Human Mouse Rat) according to manufacturer protocol (TruSeq Stranded Total RNA Sample Prep Guide, Part #15031048 Rev. E). The paired-end sequencing was performed in the Illumina NovaSeq 6000 System at Macrogen, Inc, South Korea and ∼100 M reads per sample raw data was obtained. Quality of the reads is checked with the software FASTQC and then trimming is performed with the software Trimmomatic^106^ setting a minimum base quality of 25 and a minimum length of 35 bp. Trimmed reads are then mapped on the reference genome using STAR^107^ and featureCounts^108^ is used to perform gene expression quantification. Lowly expressed genes are removed using the algorithm HTSFilter^109^ and finally differentially expressed genes are identified using edgeR^110^ setting 0.05 as the threshold of FDR significance. The RNA-Seq data analysis was done using artificial intelligence RNA-Seq (A.I.R.), Sequentia Biotech SL, Barcelona. The genes showed LogFC more than 1 for upregulated and less than -1 for downregulated DEGs were used for GO enrichment analysis in ShinyGO 0.80.^111^

### Co-immunoprecipitation

The cells were washed with ice cold PBS and collected. The total cellular extracts were prepared form these cells using RIPA lysis buffer. Total 500 μg of cellular extract was precleared with protein A-agarose beads and was incubated with anti-vinculin (Abcam) antibody overnight at 4°C in a rotator. The next day protein A-agarose beads were added and incubated for 3-4 hr at 4°C to bring down the immune complex. The beads were washed with four times with cold PBS. This immune complex was further analyzed by western blot analysis.

### Reproducibility

All functional experiments were verified at least three times independently and consistent results were obtained.

## QUANTIFICATION AND STATISTICAL ANALYSIS

Unpaired two-tailed student’s t test analyses was performed to ascertain statistical significance of comparing two mean value. Replicated samples were mentioned in figure legends as ‘n=’. Data were shown as mean value ±SEM, p value * ≤0.05; **≤0.01; ***≤0.001; ****≤0.0001. and p < 0.05 was considered to be significant.

## KEY RESOURCES TABLE

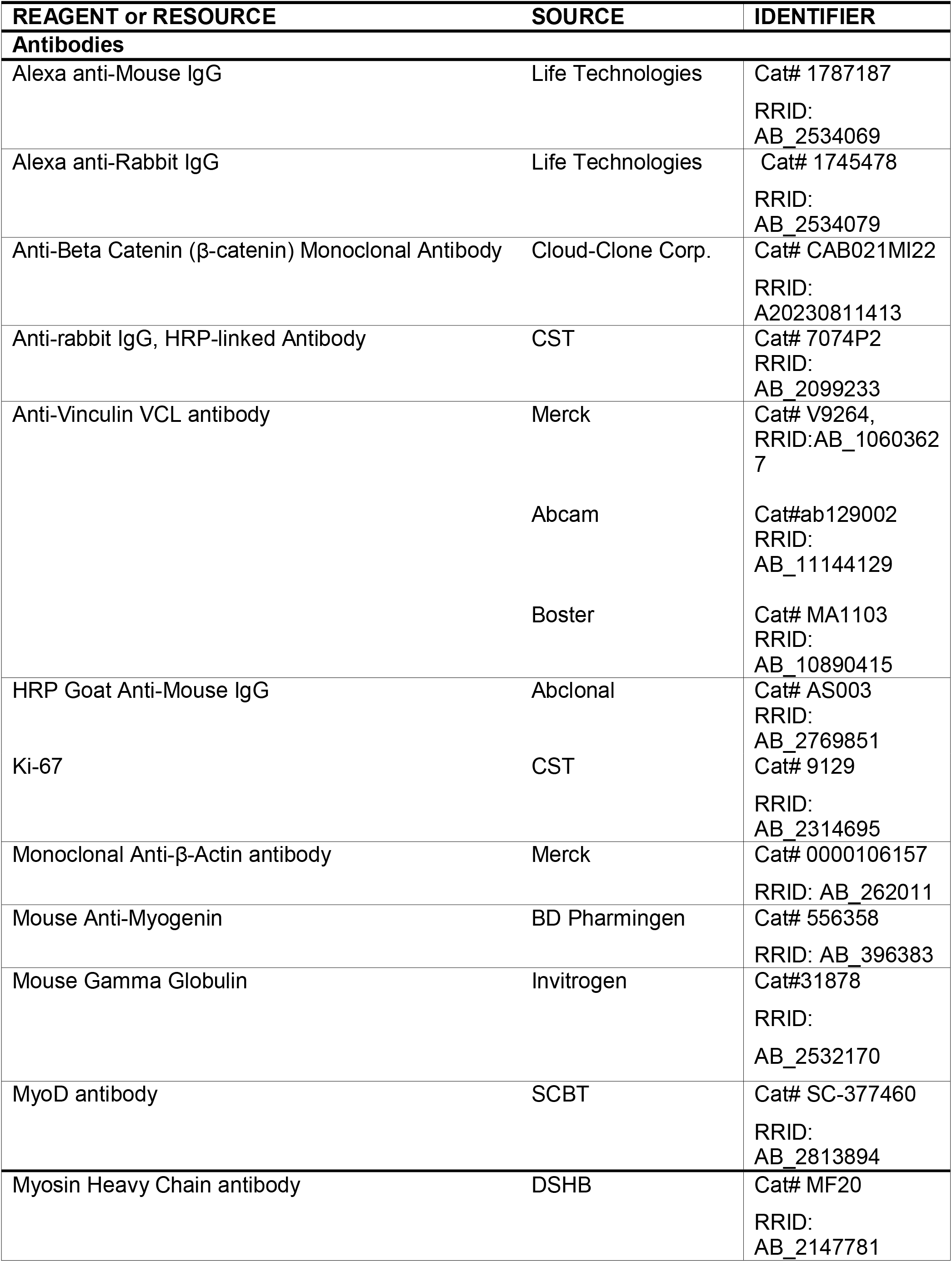
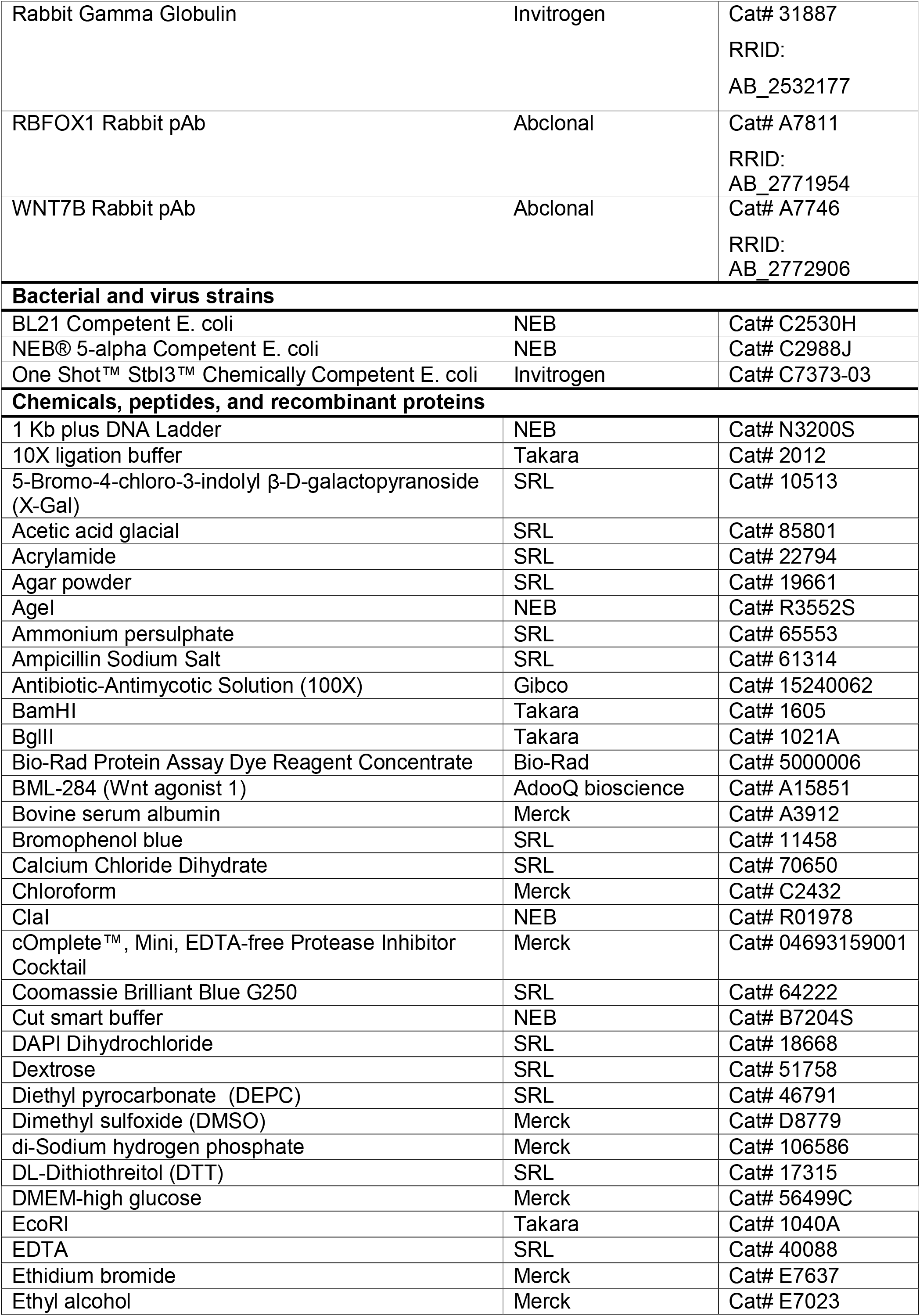
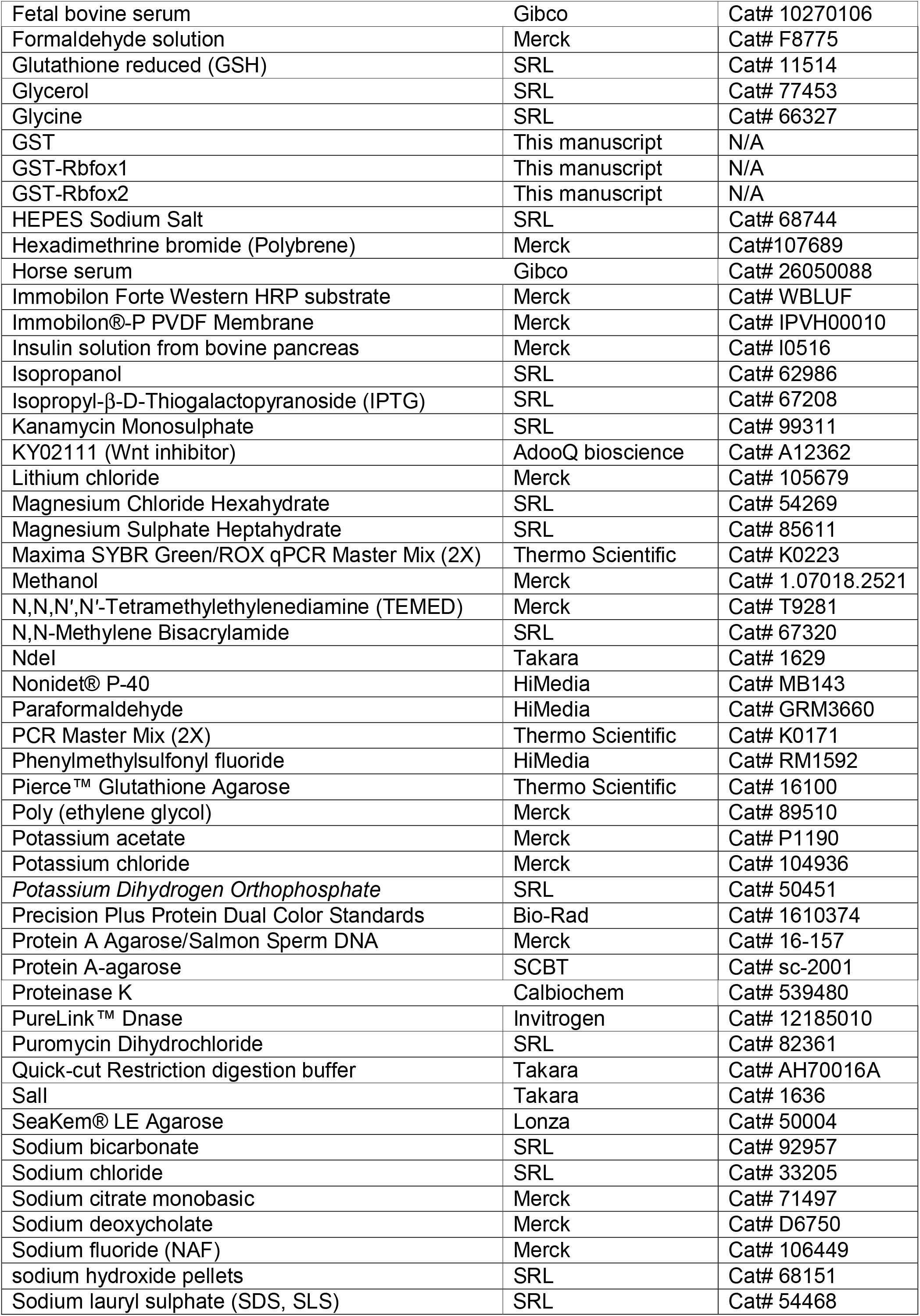
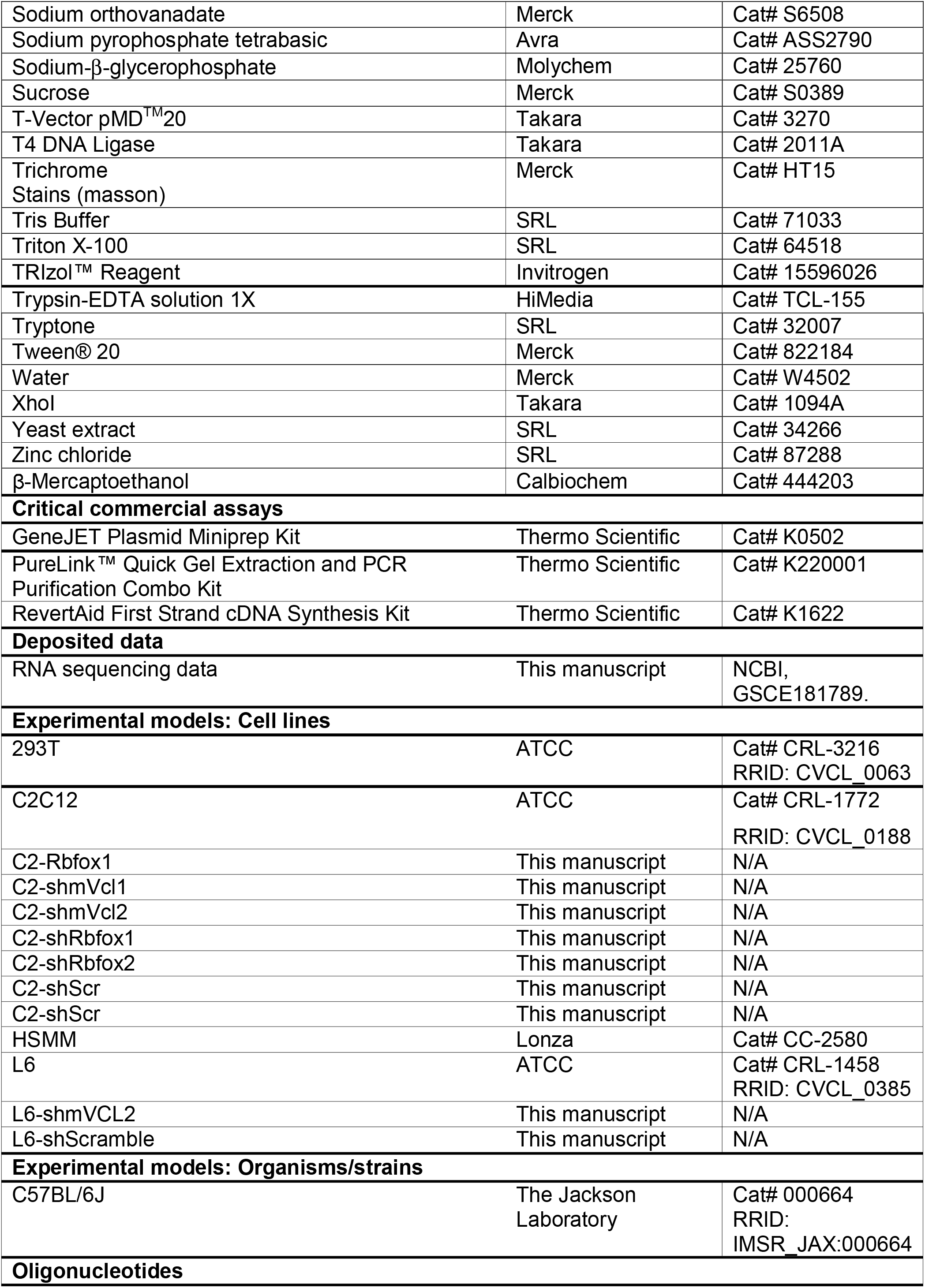
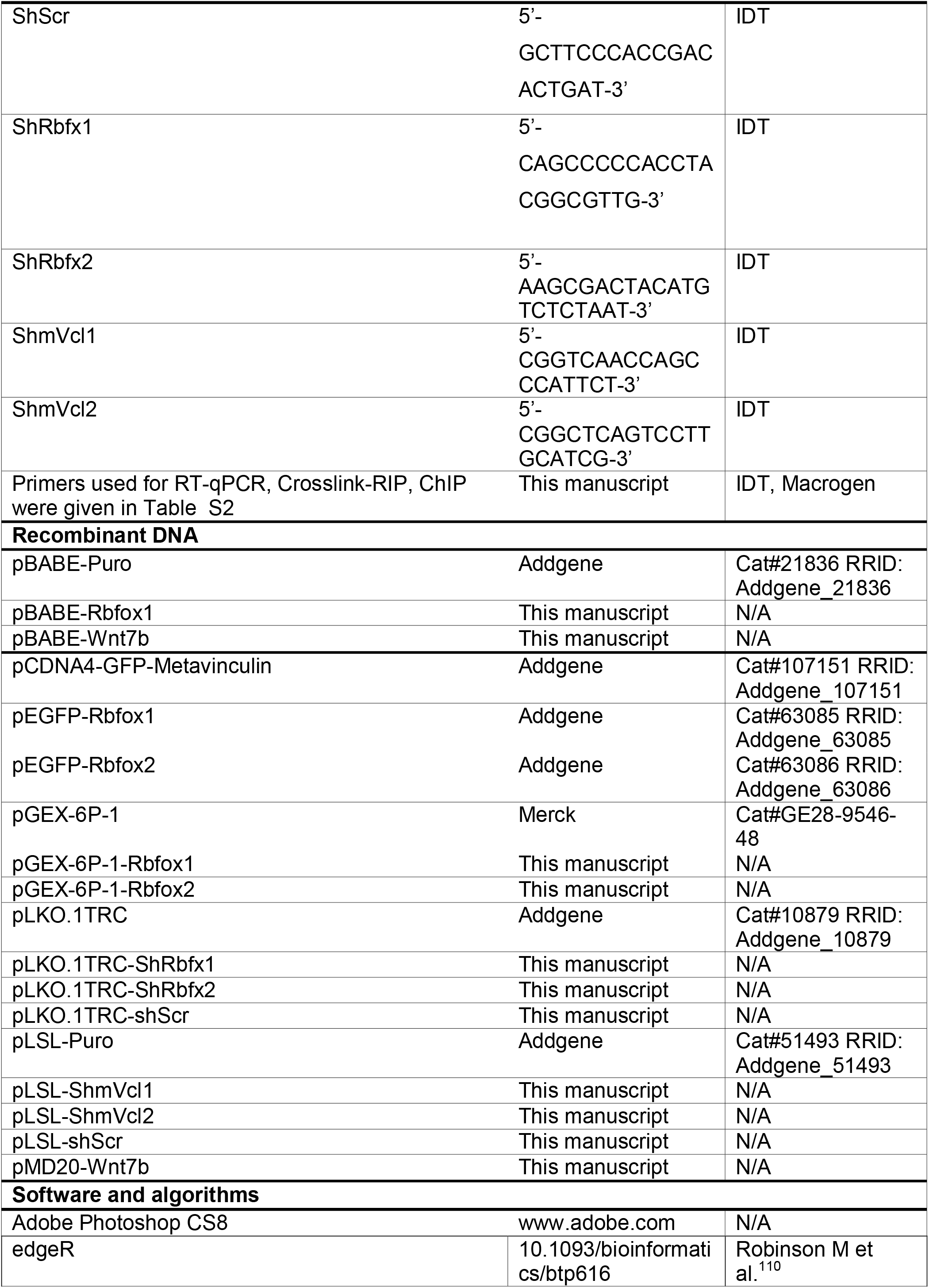
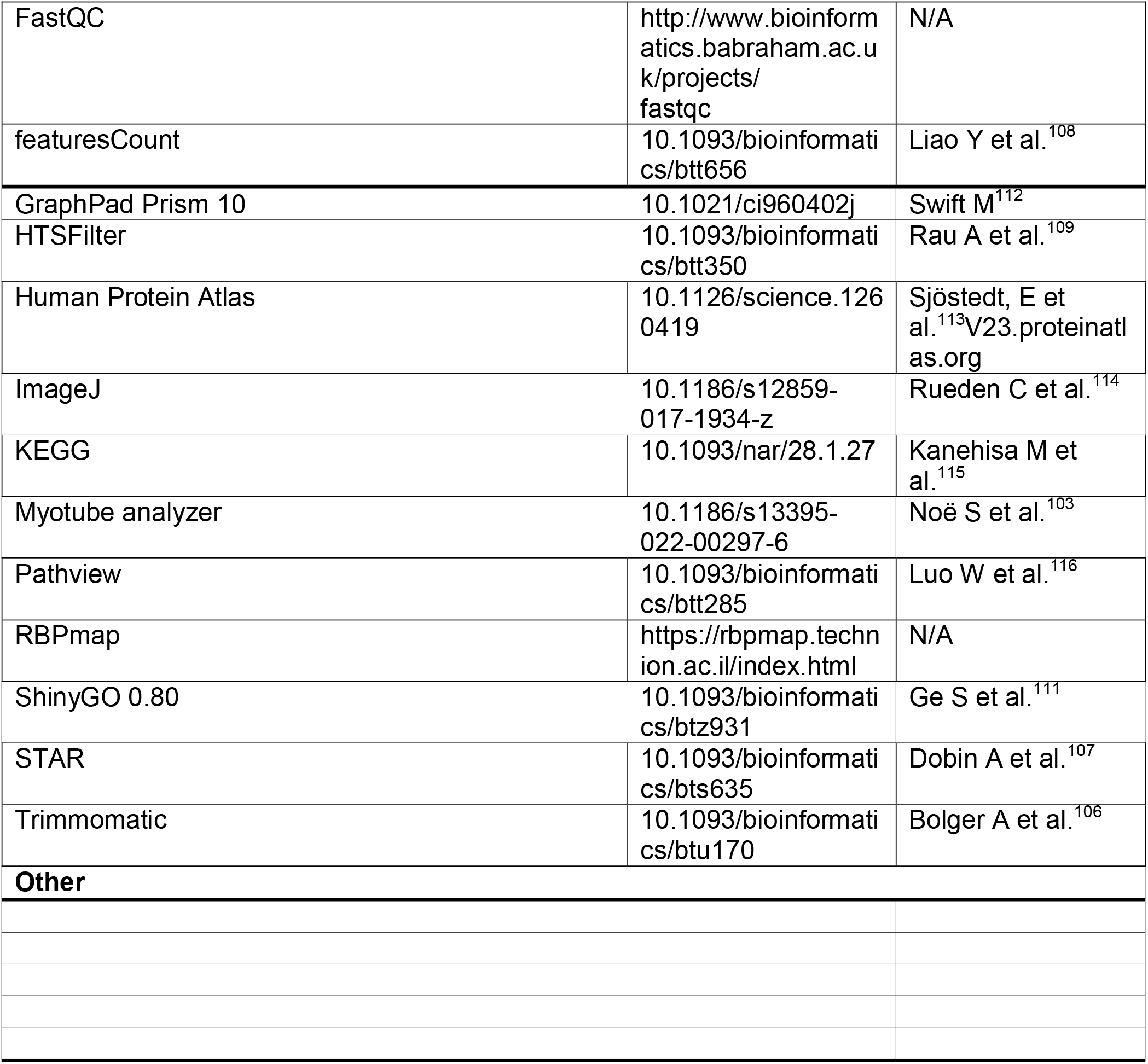

## SUPPLEMENTAL INFORMATION

**Document S1. Tables S1, S2, Figure Legends S1-S5**

**Table S3. Dataset RNA-Seq C2-shScr vs. C2-shmVcl2, related to Figure 4**

