## Supplementary Table S1, Table S2, Figure legends S1-S5 and Figures S1-S5 for "Metavinculin mediates differentiation cues into myogenic gene expression"

[3]Lead contact

**Table S1: Oligonucleotides/primers**

| S. No. | Gene | Sequence |
|---|---|---|
| **Primers used in RT-qPCR** | | |
| 1. | mVcl | F: 5'-CGCTCAGGTGGGTATAGGAG-3'<br>R: 5'-GCCATCAGCAGAGCCATG-3' |
| 2. | Vcl | F: 5'-CACCGTGTGATGTTGGTGAA-3'<br>R: 5'-ATGGCTTCAGTGTCCTTGCT-3' |
| 3. | Myh | F: 5'-CGCATCAAGGAGCTCACCTA-3'<br>R: 5'-TACTCCTCATTCAGGCCCTTG-3' |
| 4. | Myog | F: 5'-GTGTGTAAGAGGAAGTCTGT-3'<br>R: 5'-GCAAATGATCTCCTGGGTTGG-3' |
| 5. | Myod1 | F: 5'-CGCCGCCTGAGCAAAGTGAA-3'<br>R: 5'-GCCGCTGTAATCCATCATGC-3' |
| 6. | Cdkn1a | F: 5'-GACTTCGTCACGGAGACGCC-3'<br>R: 5'-GGAGTGATAGAAATCTGTCAGGC-3' |
| 7. | Actb | F: 5'-TCCATCATGAAGTGTGACGT-3'<br>R: 5'-TGCTTGCTGATCCACATC-3' |
| 8. | Rbfox1 | F: 5'-GCACCGTGTACAACACCTTC-3'<br>R: 5'-GCAAAAGCATTCATGGCACC-3' |
| 9. | Rbfox2 | F: 5'-AGGGCCGTAAAATCGAGGTG-3'<br>R: 5'-TGCAGTTGGGTAAGGGAAGC-3' |
| 10. | Rbfox3 | F: 5'-TGGCATGACCCTGTACACAC-3' |

| | | |
|---|---|---|
| | | R: 5’-CGAATTGCCCGAACATTTGC-3’ |
| 11. | Mef2dα1 | F: 5’-GGAAGTTTGGACTGATGAAG-3’<br>R: 5’-TGGCTCGTTGTACTCGGTGT-3’ |
| 12. | Mef2dα2 | F: 5'-CCTGAGGAAGAAGGGTTT-3’<br>R: 5’-GGCCCGCCGGTACTTGTC-3’ |
| 13. | Wnt7b | F: 5’-GTTTCTCTGCTTTGGCGTCCT-3’<br>R: 5’-ATCACAATGATGGCATCGGG-3’ |
| 14. | Car3 | F: 5’-ATTGCCAAAGGGGACAACCA-3’<br>R: 5’-ACCACCCCTCAGCATAGACC-3’ |
| 15. | Ccn2 | F: 5’-CTCCGTCGCAGGTCCCATC-3’<br>R: 5’-CACGCTCCGTACACAGTTCTC-3’ |
| 16. | Tagln | F: 5’-GATCGAAGCCAGTGAAGGTG-3’<br>R: 5’-TGCTGCCATATCCTTACCTTCA-3’ |
| 17. | Actg2 | F: 5’-GTTCTGGATTCGGGGGATGG-3’<br>R: 5’-TCTGTGAGATCCCGTCCAGCC-3’ |
| 18. | Myl9 | F: 5’-TGGAGGGCATGATGAACGAG-3’<br>R: 5’-CCTCCTCATCAAAGCAGGCA-3’ |
| 19. | Klhl13 | F: 5’-TGTTTCTCATATCTGGGGTCACTT-3’<br>R: 5’-GGACAAAGGCAAGGCGCTC-3’ |
| 20. | Heatr9 | F: 5’-CCCTCAATGAGCAACGCAAG-3’<br>R: 5’-ACAACCCTGGTCCCTCCTTA-3’ |
| 21. | Cxcl10 | F: 5’-ATGACGGGCCAGTGAGAATG-3’<br>R: 5’-AGGAGCCCTTTTAGACCTTTTT-3’ |
| 22. | Ccl5 | F: 5’-TTTGCCTACCTCTCCCTCGC-3’<br>R: 5’-GTTCCTTCGAGTGACAAACACGAC-3’ |
| 23. | Slurp1 | F: 5’-GCCCACGGCCATTAACTCAT-3’<br>R: 5’-ATGGGACTGTGGTTGAAGGG-3’ |
| 24. | Myc | F: 5’-CCTGTACCTCGTCCGATTCC-3’<br>R: 5’-CTTCTTGCTCTTCTTCAGAGTCG-3’ |
| 25. | Ccnd1 | F: 5’-AAAATGCCAGAGGCGGATGA-3’<br>R: 5’-CAGGGCCTTGACCGGG-3’ |
| 26. | Ascl2 | F: 5’-TGTTAACACCCGCTACTCCG-3’<br>R: 5’-GCCCCCTAACCAACTGGAAA-3’ |
| 27. | Acod1 | F: 5’-TGAAAGTGAACCACCTGACA-3’<br>R: 5’-AATCCATGGAGTGAACAGCA-3’ |
| 28. | Id2 | F: 5’-AAAAACAGCCTGTCGGACCA-3’<br>R: 5’-TGATGTCCGTGTTCAGGGTG-3’ |

| | | |
|---|---|---|
| 29. | Tcf4 | F: 5’-CATCACCAACAGCGAATGGC-3’<br>R: 5’-CTTGGATGGCCTCCAGTTCC-3’ |
| 30. | Mmp9 | F: 5’-CCAGCTGGCAGAGGCATA-3’<br>R: 5’-CACAGCGTGGTGTTCGAATG-3’ |
| **Primers used in GST-pulldown assay, crosslink-RIP and ChIP** | | |
| 31. | 5’-mVcl-Intron | F: 5’GGTAATGGCTACTGGGTTGT-3’<br>R: 5’-CTTTGTCGATGGCGGGAA-3’ |
| 32. | 3’-mVcl-Intron | F: 5’-CAGCTCAAGTTAGGTGACAGAG-3’<br>R: 5’-TGGCAGTTTATCTGAAATTTGGC-3’ |
| 33. | Myh-promoter | F: 5’-CACCCAAGCCGGGAGAAACAGCC-3’<br>R: 5'-GAGGAAGGACAGGACAGAGGCACC-3’ |
| 34. | Myog-promoter | F: 5’-GAATCACATGTAATCCACTGGA-3’<br>R: 5’-ACGCCAACTGCTGGGTGCCA-3’ |
| **Primers used for molecular cloning** | | |
| 35. | Wnt7b-cDNA | F: 5’-CGGGATCCATGCACAGAAACTTTCGAAAGTG-3’<br>R: 5’-CGAATTCCTCACTTGCAGGTGAAGACCTC-3’ |

Table S2: **Tau specificity score**

| S.No | Gene | Tissue specificity | Tissue specificity score |
|---|---|---|---|
| 1 | CELF1 | Low tissue specificity | 0.21 |
| 2 | MBNL1 | Low tissue specificity | 0.27 |
| 3 | RBFOX1 | Brain, skeletal muscle, tongue | 0.82 |
| 4 | RBFOX2 | Low tissue specificity | 0.28 |
| 5 | RBFOX3 | Brain, intestine, urinary bladder | 0.78 |
| 6 | PTB1 | Low tissue specificity | 0.13 |

Data credit: Human Protein Atlas
Data available from: V23.proteinatlas.org

### SUPPLEMENTARY FIGURE LEGENDS

**Figure S1: Myogenic gene expression during skeletal muscle differentiation, related to Figure 1, Table S1**

(A) Expression of growth arrest Cdkn1a and myogenic genes Myod1, Myog and Myh expression in C2C12 (Mouse), L6 (Rat) and HSMM (Human) myoblasts grown under growth (GM) and differentiation (DM) media.

(B) Immunofluorescence Myh staining (green) and DAPI stained (blue) nuclei showing myotube formation in various myoblasts grown under differentiation conditions and the fusion index shown in a bar graph.

Data are shown as mean value $\pm$SEM, n=3. p value **≤0.01; ***≤0.001; ****≤0.0001.

**Figure S2: Effect of Rbfox depletion on metavinculin expression and skeletal muscle differentiation, related to Figure 2, Table S1**

(A) Image showing potential Rbfoxs binding in the upstream and downstream flanking introns (purple) to the metavinculin-specific exon (red).

(B) Coomassie stain showing recombinant GST, GST-Rbfox1 and GST-Rbfox2 proteins.

(C) Bar graph showing expression of myogenic differentiation associated genes Cdkn1a, Myod1, Myog and Myh in Rbfox depleted cells.

(D) Western blot images showing metavinculin and myogenic proteins expression in Rbfox-depleted C2-shRbfx1 and C2-shRbfx2 and scramble control C2-Scr cells cultured in indicated conditions. Actb act as a loading control.

(E and F) Representative Myh (green) stain and counter stain DAPI (blue) images showing myotube formation (E) and quantified fusion index in a graph (F) in Rbfox depleted and control cells grown under DM media.

(G) Relative expression of Rbfox1 known target Mef2d$\alpha$ in C2C12 cells contains ectopic Rbfox1 or vector.

Data are shown as mean value $\pm$SEM, n=3. p value * ≤0.05; **≤0.01; ****≤0.0001; ns-not significant.

**Figure S3: Metavinculin depletion inhibits MyoD occupancy at Myog and Myh promoters, related to Figure 3, Table S1**

(A and B) Inhibition of MyoD binding to Myog (A) and Myh (B) promoters in metavinculin depleted C2-shmVcl2 cells is depicted in a bar graph.

Data are shown as mean value ±SEM, n=3. p value ***≤0.001; ****≤0.0001.

**Figure S4: DEGs in metavinculin depleted cells, related to Figure 4**

(A) Cellular Components GO terms of DEGs in metavinculin depleted C2C12 cells are shown in a bar plot.

(B) Image showing upregulated DEGs (highlighted in red) involved in DNA replication (Eukaryotes) KEGG pathway.

(C) FPKM values of cyclins Ccna2, Ccnb2, Ccbe2 and Ccnf in C2-shmVcl2 and C2-Scr cells.

(D) Bar plot showing downregulated DEGs enrichment in KEGG pathways.

Data are shown as mean value ±SEM, n=3. p value *≤0.05; **≤0.01.

**Figure S5: Downregulated DEGs in metavinculin depleted cells on KEGG Wnt pathway, related to Figure 4**

(A) Network image showing downregulated DEGs (highlighted in red) involved in KEGG Wnt pathway.

(B) FPKM values of non-canonical Wnt7b and Wnt11 in C2-shmVcl2 and C2-Scr cells. Data are shown as mean value ±SEM, n=3. p value ***≤0.001; ns-not significant.

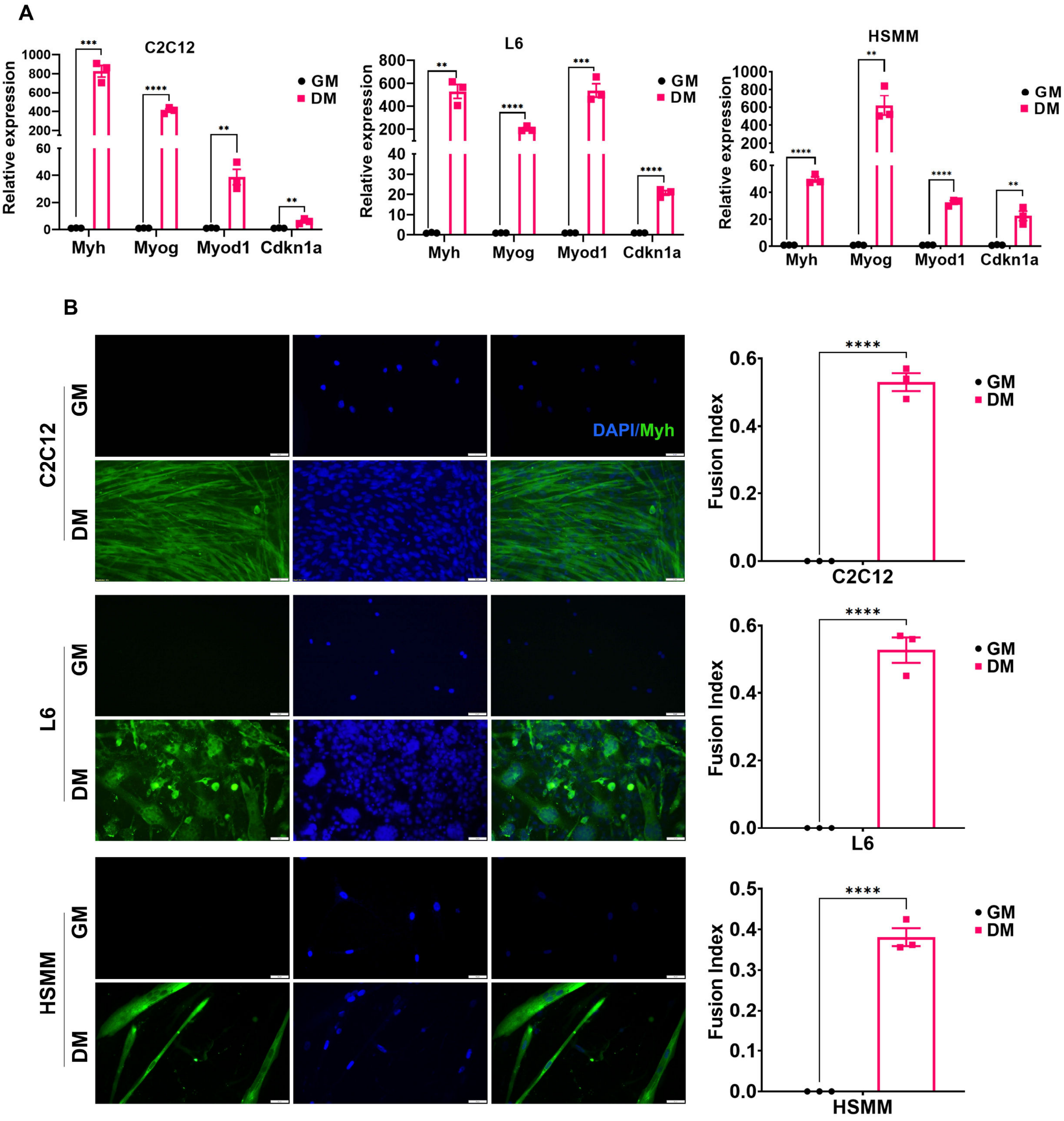
A
C2C12
L6
HSMM
Relative expression
GM
DM
Myh
Myog
Myod1
Cdkn1a
B
C2C12
L6
HSMM
DAPI/Myh
Fusion Index


Figure S1

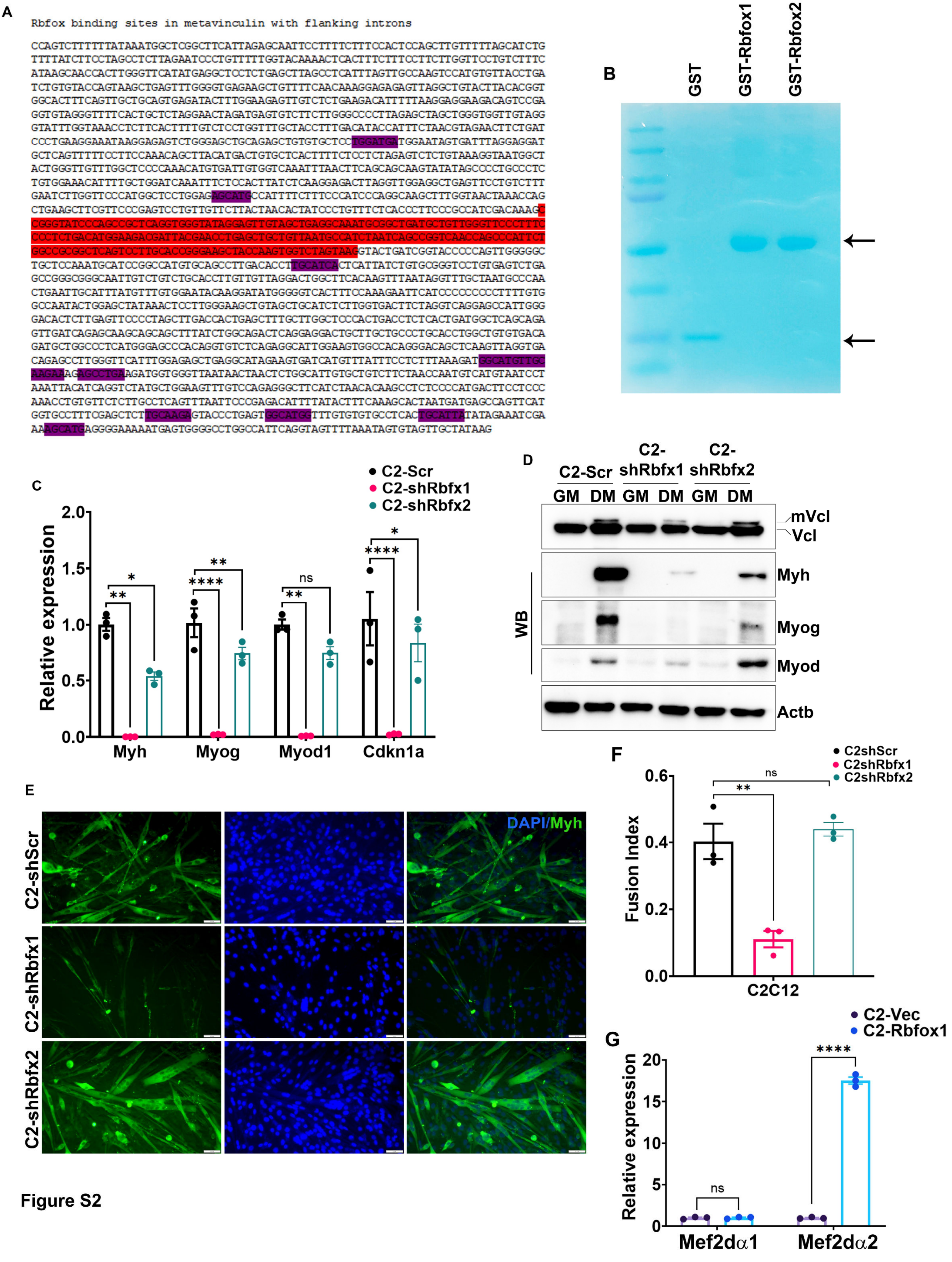
A
Rbfox binding sites in metavinculin with flanking introns
CCAGTCTTTTTTATAAATGGCTCGGCTTCATTAGAGCAATTCCTTTTCTTTCCACTCCAGCTTGTTTTTAGCATCTG
TTTTATCTTCCTAGCCTCTTAGAATCCCTGTTTTTGGTACAAAACTCACTTTCTTTCCTTCTTGGTTCCTGTCTTTC
ATAAGCAACCACTTGGGTTCATATGAGGCTCCTCTGAGCTTAGCCTCATTTAGTTGCCAAGTCCATGTGTTACCTGA
TCTGTGTACCAGTAAGCTGAGTTTGGGGTGAGAAGCTGTTTTCAACAAAGGAGAGAGTTAGGCTGTACTTACACGGT
GGCACTTTCAGTTGCTGCAGTGAGATACTTTGGAAGAGTTGTCTCTGAAGACATTTTTAAGGAGGAAGACAGTCCGA
GGTGTAGGGTTTTCACTGCTCTAGGAACTAGATGAGTGTCTTCTTGGGCCCCTTAGAGCTAGCTGGGTGGTTGTAGG
GTATTTGGTAAACCTCTTCACTTTTGTCTCCTGGTTTGCTACCTTTGACATACCATTTCTAACGTAGAACTTCTGAT
CCCTGAAGGAAATAAGGAGAGTCTGGGAGCTGCAGAGCTGTGTGCTCCTGGATGATGGAATAGTGATTTAGGAGGAT
GCTCAGTTTTTCCTTCCAAACAGCTTACATGACTGTGCTCACTTTTCTCCTCTAGAGTCTCTGTAAAGGTAATGGCT
ACTGGGTTGTTTGGCTCCCCAAACATGTGATTGTGGTCAAATTTAACTTCAGCAGCAAGTATATAGCCCCTGCCCTC
TGTGGAAACATTTTGCTGGATCAAATTTCTCCACTTATCTCAAGGAGACTTAGGTTGGAGGCTGAGTTCCTGTCTTT
GAATCTTGGTTCCCATGGCTCCTGGAGAGCATGCCATTTTCTTTCCCATCCCAGGCAAGCTTTGGTAACTAAACCAG
CTGAAGCTTCGTTCCCGAGTCCTGTTGTTCTTACTAACACTATCCCTGTTTCTCACCCTTCCCGCCATCGACAAAGC
CGGGTATCCCAGCCGCTCAGGTGGGTATAGGAGTTGTAGCTGAGGCAAATGCGGCTGATGCTGTTGGGTTCCCTTTC
CCCTCTGACATGGAAGACGATTACGAACCTGAGCTGCTGTTAATGCCATCTAATCAGCCGGTCAACCAGCCCATTCT
GGCCGCGGCTCAGTCCTTGCACCGGGAAGCTACCAAGTGGTCTAGTAAGGTACTGATCGGTACCCCCAGTTGGGGGC
TGCTCCAAATGCATCCGGCCATGTGCAGCCTTGACACCTTGCATCACTCATTATCTGTGCGGGTCCTGTGAGTCTGA
GCCGGGCGGGCAATTGTCTGTCTGCACCTTGTTGTTAGGACTGGCTTCACAAGTTTAATAGGTTTGCTAATGCCCAA
CTGAATTGCATTTATGTTTGTGGAATACAAGGATATGGGGGTCACTTTCCAAAGAATTCATCCCCCCCCCTTTTGTG
GCCAATACTGGAGCTATAAACTCCTTGGGAAGCTGTAGCTGCATCTCTTGGTGACTTCTAGGTCAGGAGCCATTGGG
GACACTCTTGAGTTCCCCTAGCTTGACCACTGAGCTTTGCTTGGCTCCCACTGACCTCTCACTGATGGCTCAGCAGA
GTTGATCAGAGCAAGCAGCAGCTTTATCTGGCAGACTCAGGAGGACTGCTTGCTGCCCTGCACCTGGCTGTGTGACA
GATGCTGGCCCTCATGGAGCCCACAGGTGTCTCAGAGGCATTGGAAGTGGCCACAGGGACAGCTCAAGTTAGGTGA
CAGAGCCTTGGGTTCATTTGGAGAGCTGAGGCATAGAAGTGATCATGTTTATTTCCTCTTTAAAGATGGCATGTTGC
AAGAAAGAGCCTGAAGATGGTGGGTTAATAACTAACTCTGGCATTGTGCTGTCTTCTAACCAATGTCATGTAATCCT
AAATTACATCAGGTCTATGCTGGAAGTTTGTCCAGAGGGCTTCATCTAACACAAGCCTCTCCCCATGACTTCCTCCC
AAACCTGTGTTCTCTTGCCTCAGTTTAATTCCCGAGACATTTTATACTTTCAAAGCACTAATGATGAGCCAGTTCAT
GGTGCCTTTCGAGCTCTTGCAAGAGTACCCTGAGTGGCATGGTTTGTGTGTGCCTCACTGCATTATATAGAAATCGA
AAAGCATGAGGGGAAAAATGAGTGGGGCCTGGCCATTCAGGTAGTTTTAAATAGTGTAGTTGCTATAAG
B
GST
GST-Rbfox1
GST-Rbfox2
C
C2-Scr
C2-shRbfx1
C2-shRbfx2
Relative expression
2.0
1.5
1.0
0.5
0.0
**
*
****
**
ns
****
*
Myh
Myog
Myod1
Cdkn1a
D
C2-Scr
C2-shRbfx1
C2-shRbfx2
GM DM GM DM GM DM
mVcl
Vcl
Myh
Myog
Myod
Actb
WB
E
C2-shScr
C2-shRbfx1
C2-shRbfx2
DAPI/Myh
F
C2shScr
C2shRbfx1
C2shRbfx2
Fusion Index
0.6
0.4
0.2
0.0
ns
**
C2C12
G
C2-Vec
C2-Rbfox1
Relative expression
20
15
10
5
0
ns
****
Mef2dα1
Mef2dα2

Figure S2

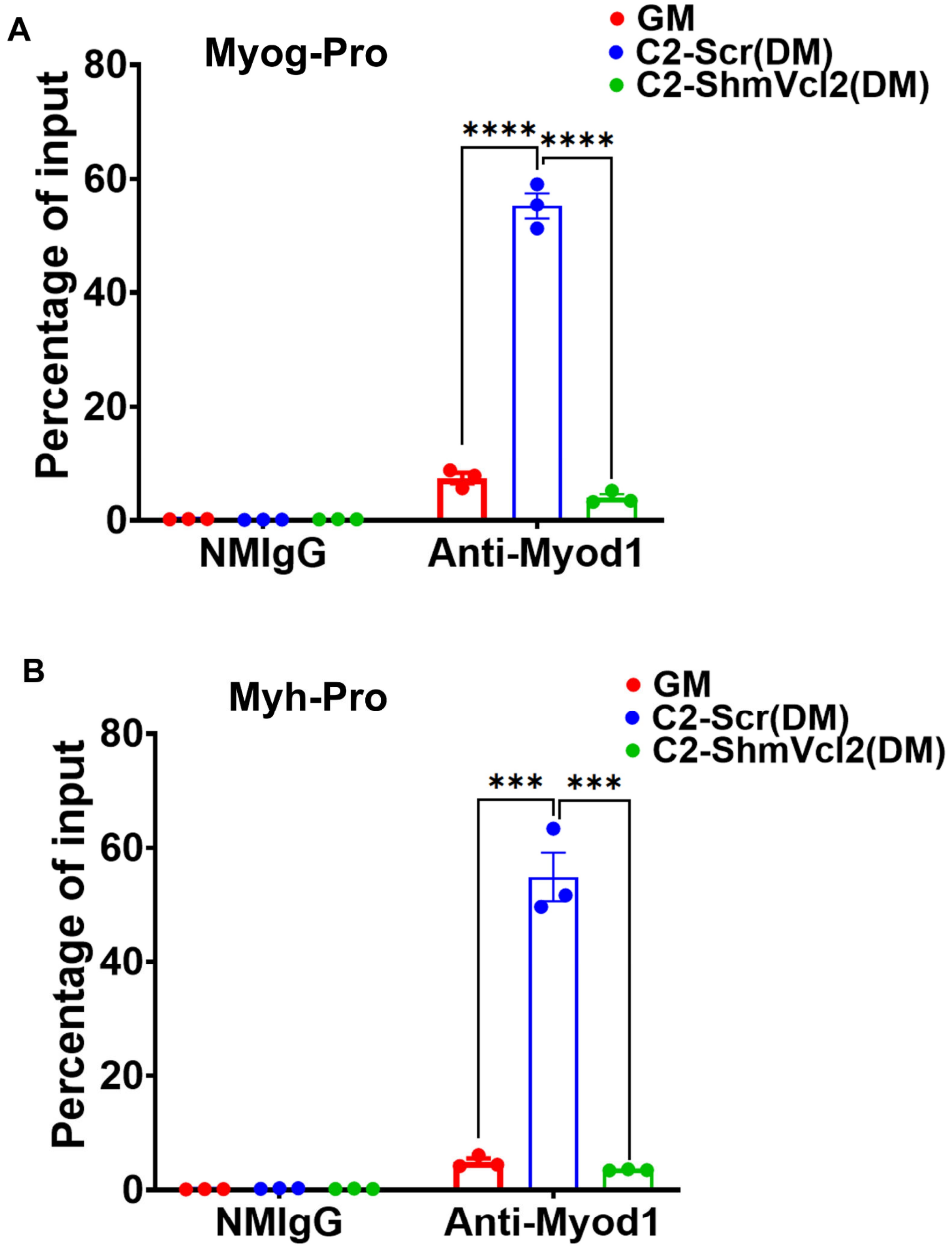
A
Myog-Pro
GM
C2-Scr(DM)
C2-ShmVcl2(DM)
Percentage of input
80
60
40
20
0
****
****
NMIgG
Anti-Myod1
B
Myh-Pro
GM
C2-Scr(DM)
C2-ShmVcl2(DM)
Percentage of input
80
60
40
20
0
***
***
NMIgG
Anti-Myod1

**Figure S3**

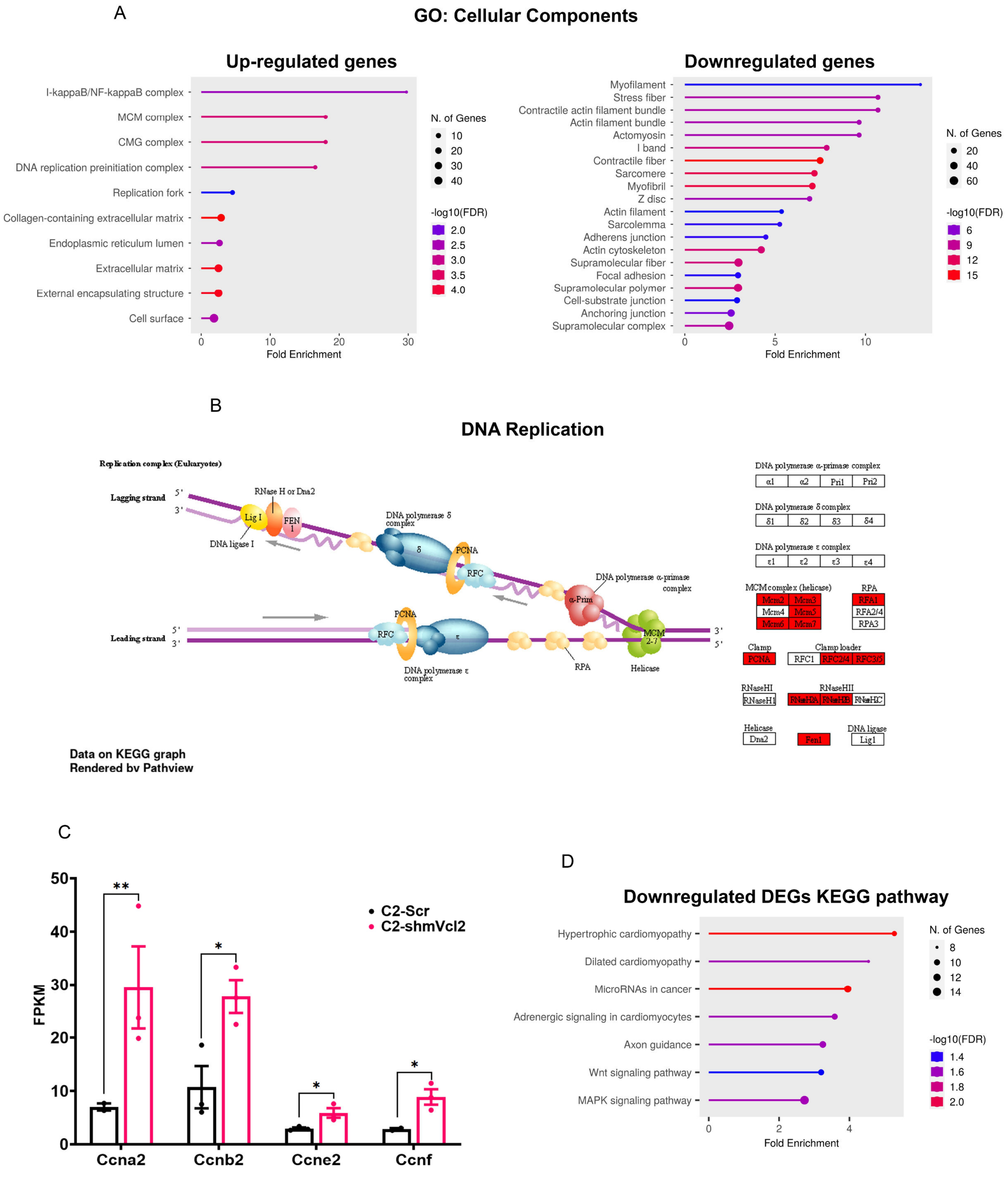
A
GO: Cellular Components
Up-regulated genes
I-kappaB/NF-kappaB complex
MCM complex
CMG complex
DNA replication preinitiation complex
Replication fork
Collagen-containing extracellular matrix
Endoplasmic reticulum lumen
Extracellular matrix
External encapsulating structure
Cell surface
0 10 20 30
Fold Enrichment
N. of Genes
10
20
30
40
-log10(FDR)
2.0
2.5
3.0
3.5
4.0
Downregulated genes
Myofilament
Stress fiber
Contractile actin filament bundle
Actin filament bundle
Actomyosin
I band
Contractile fiber
Sarcomere
Myofibril
Z disc
Actin filament
Sarcolemma
Adherens junction
Actin cytoskeleton
Supramolecular fiber
Focal adhesion
Supramolecular polymer
Cell-substrate junction
Anchoring junction
Supramolecular complex
0 5 10
Fold Enrichment
N. of Genes
20
40
60
-log10(FDR)
6
9
12
15
B
DNA Replication
Replication complex (Eukaryotes)
Lagging strand
Leading strand
RNase H or Dna2
Lig I
FEN 1
DNA ligase I
DNA polymerase δ complex
PCNA
RFC
DNA polymerase α-primase complex
α-Prim
MCM 2-7
RPA
Helicase
DNA polymerase ε complex
DNA polymerase α-primase complex
α1 α2 Pri1 Pri2
DNA polymerase δ complex
δ1 δ2 δ3 δ4
DNA polymerase ε complex
ε1 ε2 ε3 ε4
MCM complex (helicase)
Mcm2 Mcm3
Mcm4 Mcm5
Mcm6 Mcm7
RPA
RFA1
RFA2/4
RPA3
Clamp
PCNA
Clamp loader
RFC1 RFC2/4 RFC3/5
RNaseHI
RNaseH1
RNaseHII
RNaseH2A RNaseH2B RNaseH2C
Helicase
Dna2
Fen1
DNA ligase
Lig1
Data on KEGG graph
Rendered by Pathview
C
FPKM
0 10 20 30 40 50
**
*
*
*
C2-Scr
C2-shmVcl2
Ccna2
Ccnb2
Ccne2
Ccnf
D
Downregulated DEGs KEGG pathway
Hypertrophic cardiomyopathy
Dilated cardiomyopathy
MicroRNAs in cancer
Adrenergic signaling in cardiomyocytes
Axon guidance
Wnt signaling pathway
MAPK signaling pathway
0 2 4
Fold Enrichment
N. of Genes
8
10
12
14
-log10(FDR)
1.4
1.6
1.8
2.0

**Figure S4**

a

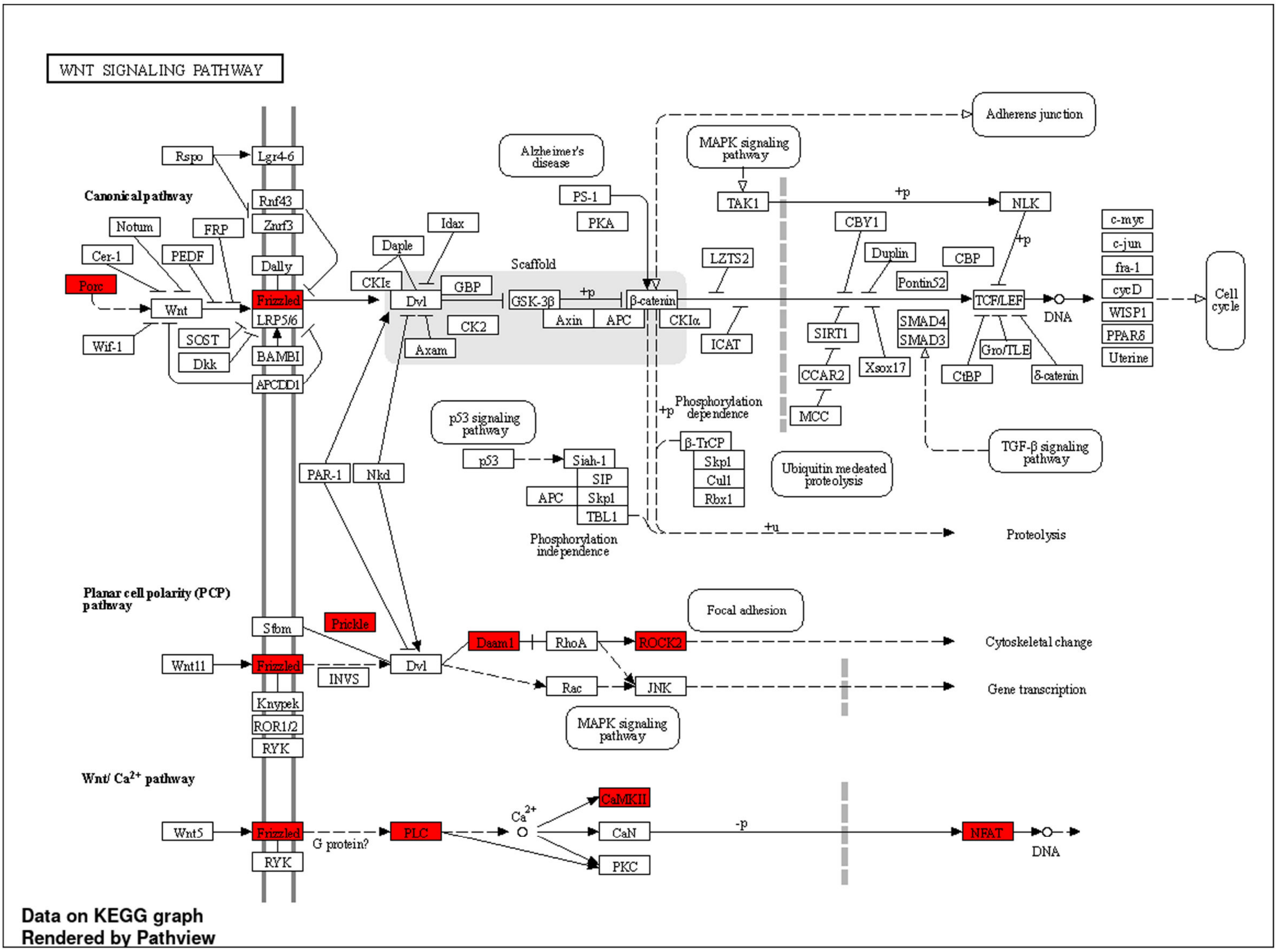
WNT SIGNALING PATHWAY
Canonical pathway
Planar cell polarity (PCP) pathway
Wnt/ Ca2+ pathway
Data on KEGG graph
Rendered by Pathview

b

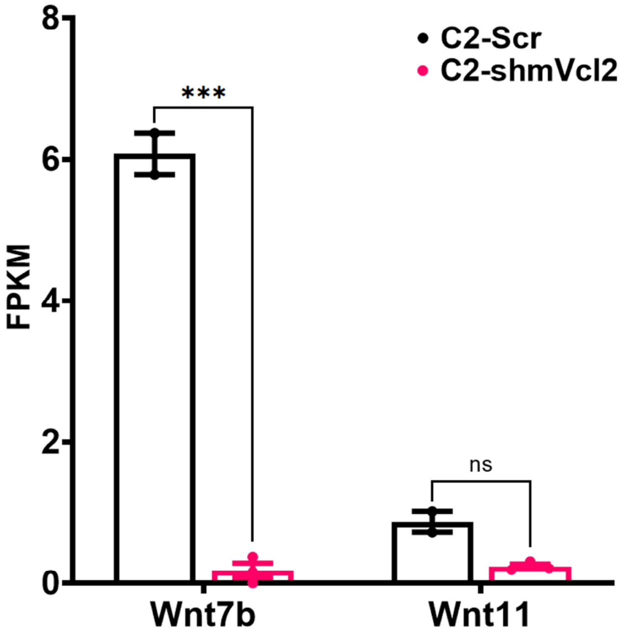
C2-Scr
C2-shmVcl2
FPKM
***
ns
Wnt7b
Wnt11

Figure S5
